# Resolution of collapsed forks is separate from completion of DNA synthesis

**DOI:** 10.1101/2025.03.03.641087

**Authors:** Sara C Conwell, Steven N Dahmen, Matthew T. Cranford, Ruthie J. Mulvaney, David T Long, David Cortez, James M. Dewar

## Abstract

Replication fork collapse at single-strand DNA breaks (SSBs) poses a serious threat to genome stability. Using *Xenopus* egg extracts, we show that a replication fork encountering an SSB on either the leading- or lagging-strand template produces a single-ended double-strand break (seDSB). These broken ends are efficiently resolved by homologous recombination to yield D-loops and erroneous end-to-end fusions. Surprisingly, DNA synthesis downstream of an seDSB is highly inefficient. In contrast, when two forks converge at an SSB, they generate a double-ended DSB (deDSB) that efficiently completes DNA synthesis through double-strand break repair that is not dependent on homologous recombination. Leading, but not lagging, seDSBs can undergo extensive nucleolytic degradation that disassembles the divergent fork. These ‘secondary collapse’ events efficiently resolve seDSBs but without completion of DNA synthesis. Moreover, PARP inhibition can enhance fork collapse at unmodified SSBs but not at abasic site SSBs, contrary to expectations. Our findings distinguish end resolution from replication completion and demonstrate flexibility in how PARP inhibition affects fork collapse.

## Introduction

Tens of thousands of single-strand DNA breaks (SSBs) arise spontaneously each day^1–3^. When a replication fork encounters an SSB, it can be converted to a double-strand break (DSB)^4–11^. This process is called “fork collapse” and blocks DNA synthesis. Fork collapse occurs every cell cycle in humans^12^ in response to SSBs generated by topoisomerases^13^, transcription^14–17^, viral infection^18^, antiviral responses^19^, lagging strand maturation^20^, abasic sites^21–24^, defective DNA repair^23^, and also genome editing^25,26^. If a collapsed fork is not properly addressed, it is highly mutagenic and potentially oncogenic^6,27–30^. Fork collapse is induced by many chemotherapeutics^31^ and is crucial to^11,32–35^ the cancer cell killing effects of PARP inhibitors^36–41^, which limit SSB repair^1,39–41^. Most knowledge of fork collapse has come from studying ‘complex’ SSBs that are chemically modified or protected by SSB-inducing enzymes^4,42^. However, many spontaneous SSBs are simple discontinuities in the phosphodiester backbone^1,20,21,23,43–45^. Whether fork collapse at these ‘simple’ SSBs involves the same mechanisms remains unclear. PARP is one factor that detects SSBs, but at least two other SSB recognition pathways exist^46–50^. Thus, the type of SSB may influence the likelihood of fork collapse by PARP inhibitors, but this remains to be tested. Although multiple pathways can resolve collapsed forks^6,9,42,51–70^, it is unclear how their usage is determined and whether additional pathways remain undiscovered. Moreover, it is unknown how DNA synthesis is completed downstream of a collapsed fork. It is imperative to address these questions given the profound implications of fork collapse for normal cellular function and human health.

When a replication fork encounters an SSB, the fork can break to form a single-ended DSB (seDSB)^5–7,51,52,56^, leaving a sister DNA duplex with a gap that is rapidly filled and ligated^7^. In parallel, the replicative helicase and associated proteins (‘replisome’) are lost, but through different mechanisms depending on which of the two parental strands contains the SSB^7^. Encounter with an SSB in the leading strand template (‘leading collapse’) causes rapid replisome dissociation^7^ due to loss of the translocating strand (Fig. S1A)^71,72^. An SSB on the lagging strand template (‘lagging collapse’) leads to replisome translocation over the downstream parental duplex, which triggers the replisome unloading pathway that normally operates during replication termination (Fig. S1B)^7,73^. If the lagging SSB is shielded by a DNA-binding protein then the replisome continues to unwind (Fig. S1C)^52^. This is thought to produce a 5’ flap that becomes double-stranded upon lagging strand synthesis to form a double-ended DSB (deDSB) behind the fork^51–53,74^. deDSBs also form in response to leading SSBs^51–53,75,76^ likely via arrival of a converging fork after seDSB formation^6,52,56^ (Fig. S1D). Fork convergence at an SSB is reported to generate a deDSB^76,77^, but an seDSB intermediate has not been observed, and extensive re-replication can occur^27,77^. Thus, there is a lack of direct evidence supporting the model that forks converge upon an SSB to generate a seDSB that is then converted to a deDSB.

DSB resolution at collapsed forks is critically dependent on homologous recombination^4,6,9,42,51–55^ and likely involves the formation of a RAD51-dependent D-loop^55,78^, regardless of whether an seDSB or a deDSB is formed^42^. In contrast, DSBs formed independently of replication undergo both homologous recombination and end-joining^52,79^. Some evidence suggests that end joining may occur following fork collapse^56,57,80–84^, but it remains unclear whether this is a major outcome. Notably, seDSBs produced by leading versus lagging collapse exhibit different DNA structures, suggesting that different mechanisms might operate^7^. Accordingly, resolution of leading seDSBs requires specific proteins^51^. However, no other mechanistic differences have been identified. Overall, it remains unclear to what extent HR-independent mechanisms act on replication-dependent DSBs, and whether unique mechanisms exist for leading versus lagging seDSBs.

Homologous recombination is crucial for resolving DSBs at collapsed forks, but the downstream events that restart or complete DNA synthesis remain uncertain. Once a D-loop forms at an seDSB, it is capable of extensive DNA synthesis^85–87^ on both strands^88–90^ through ‘break induced replication’ (BIR), which is well established in eukaryotes^6,55–65^ and may function analogously to bacterial replication restart pathways^91–95^. However, several observations call into question the role of BIR as a major mechanism of replication restart. First, BIR is detected through error-prone repair events^58,65^, which may not represent most fork collapse outcomes. Second, replication restart involves key proteins that are not conserved in eukaryotes^51^. Third, DNA synthesis by BIR is strongly negatively regulated^56^. Fourth, fork collapse in yeast mostly results in deDSBs^51,56^. Finally, human BIR proteins^96,97^ are dispensable for resolving DNA ends following fork collapse^52^. Thus, BIR may be rare, as for other BIR-like pathways^66–70^. However, it is complicated to analyze BIR because interfering with seDSB resolution results in deDSB formation by fork convergence^52^. Hence, it is unclear how effectively collapsed forks complete DNA synthesis, with or without fork convergence to generate a deDSB.

To understand the causes and consequences of fork collapse, we replicated ‘simple’ SSBs using *Xenopus* egg extracts^98,99^. These extracts mimic the nuclear proteome of human cells^100,101^, support many DNA repair pathways^55,102–106^, and recapitulate fork collapse mechanisms that function in human cells^7,52^. This approach allowed us to induce fork collapse so we could monitor the resulting DNA structures formed and assess whether DNA synthesis restarted. We found that either leading or lagging collapse induced seDSBs that were efficiently resolved by RAD51 to generate D-loops, but also erroneous end-fusions. Surprisingly, restart of DNA synthesis was not detected. In contrast, when forks converged upon an SSB, the resulting deDSBs were efficiently resolved in a partially-RAD51 dependent manner that efficiently completed DNA synthesis. We observed similar outcomes at SSBs generated by Cas9 nickases. Unexpectedly, seDSBs generated by leading, but not lagging, template SSBs could undergo extensive nucleolytic degradation that disassembled the divergent fork. These ‘secondary collapse’ events efficiently resolved DNA ends but did not restart or complete DNA synthesis. Finally, we found that fork collapse at ‘simple’ SSBs could be exacerbated by PARP inhibition, while collapse at abasic sites could be resistant, indicating that certain SSBs could be preferentially repaired by PARP. Overall, our data establish resolution of broken ends as mechanistically distinct from restart or completion of DNA synthesis and reveal that the effects of PARP inhibition on fork collapse are highly flexible.

## Results

### Leading collapse elicits DSB resolution but not replication restart

We examined the events following replication of a Single Strand Break (SSB) on the leading strand template (‘leading collapse’). To this end, we used *Xenopus* egg extracts (‘extracts’) to replicate plasmid DNA containing site- and strand-specific breaks, as previously described^7,107^ (Fig. 1A). We used nicking enzymes to cleave the phosphodiester backbone, which generated a plasmid template containing a series of site- and strand-specific ‘simple’ SSBs that could not be regenerated because the DNA was purified from the nicking enzymes. Each SSB was flanked by *tet* operator (*tetO*) sequences, bound by *tet* repressor protein (TetR), to prevent rapid religation of SSBs in extracts (Fig. 1A) without affecting DNA replication^7,107^. SSBs were flanked by a *lac* operator (*lacO*) array that could be bound by *lac* repressor (LacR) to block converging forks^98^ and thus ensure either leading or lagging collapse. TetR does not completely block SSB repair, so the three tandem SSBs we used previously resulted in some molecules that were religated and did not collapse^7,107^. We therefore increased the number of SSBs to five in this study (Fig. S1E). We also incorporated an internal control plasmid that had previously been replicated (Fig. S1F) to ensure precise measurements of fork collapse.

**Figure 1:**
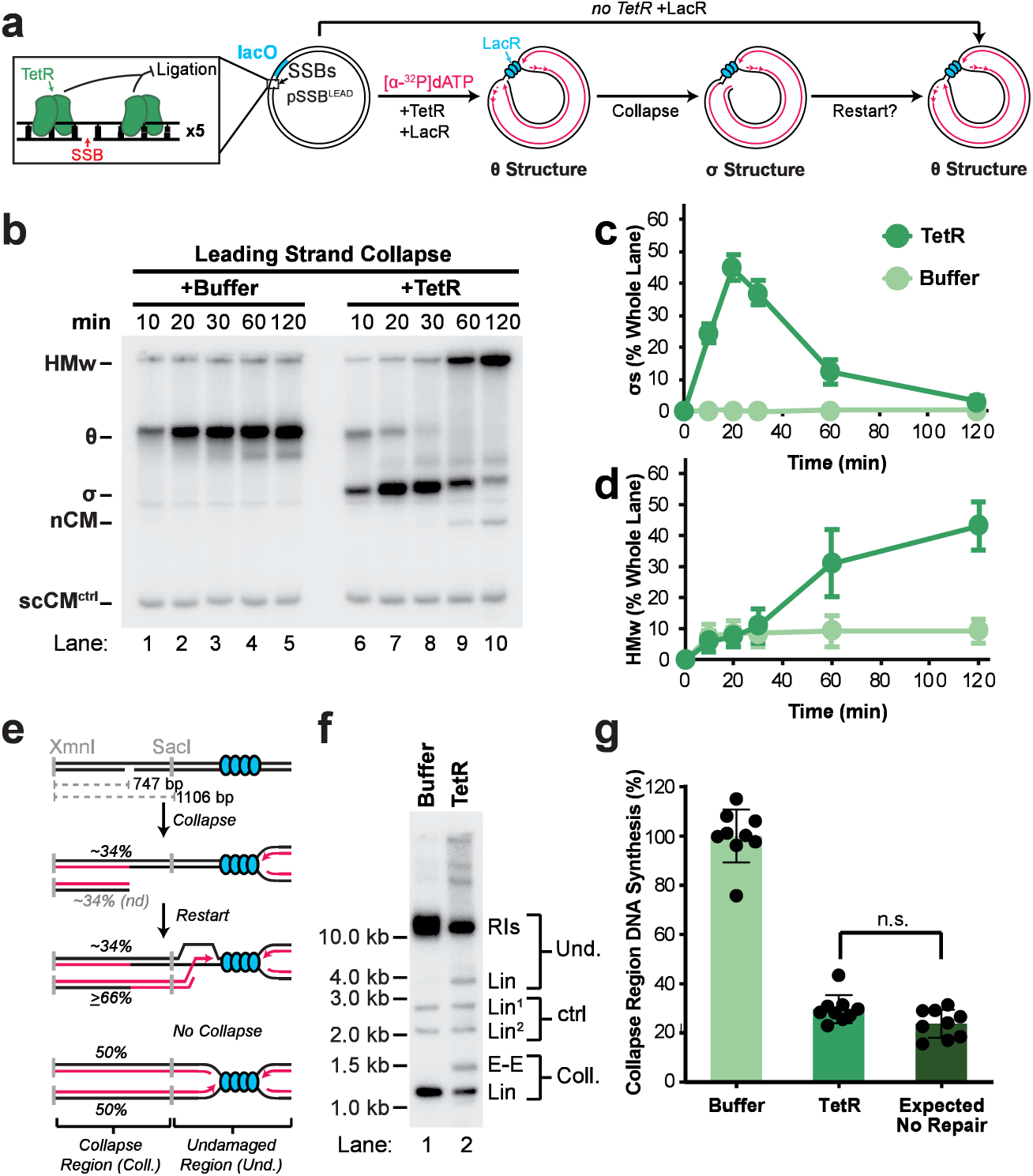
Leading seDSBs are resolved but do not restart DNA synthesis. **(A)** pSSB^LEAD^ was replicated using *Xenopus* egg extracts in the presence of (+TetR) to stabilize the SSBs. TetR was omitted in the buffer control (+Buffer), which allowed religation of SSBs prior to replication and prevented collapse (as in^7^). LacR was included in the reaction to impede converging replication forks and ensure strand-specific fork collapse at a single fork. Nascent strands were radiolabeled by inclusion of [α-^32^P] dATP. Fully replicated plasmid DNA (scCM^ctrl^) served as a loading control. See also Fig. S1F. **(B)** Samples from (a) were separated on an agarose gel and visualized by autoradiography. **(C)** Quantification of σ structures (σs) from (b). Mean ± S.D., n = 9 independent experiments. **(D)** Quantification of high molecular weight products (HMw) from (b). Mean ± S.D., n = 9 independent experiments. **(E)** Cartoon depicting the assay for restart of DNA synthesis in the collapse region. See also Fig. S1M. **(F)** Purified DNA samples from T=120 in (b) were digested with XmnI and SacI, then separated on an agarose gel and visualized by autoradiography. **(G)** Quantification of collapse region DNA synthesis from (f) as in (e). Signal was normalized to control fragment ‘Lin^1^’. Expected values for no repair account for replication efficiency and collapse efficiency. See also Fig. S1M. Mean ± S.D., n = 9 independent experiments. Data were analyzed by one-way analysis of variance (ANOVA) and Dunnett’s multiple-comparison method.

We first tested whether our revised approach more efficiently induced fork collapse. pSSB^LEAD^ was incubated with TetR and LacR, then replicated in extracts (Fig. 1B). Without TetR, SSBs were religated (as in ^7^), which produced θ structures (Fig. 1B, lanes 1-5) corresponding to converging forks at the LacR array^98^. Including TetR stabilized SSBs and generated σ structures (σ; Fig. 1B, lanes 6-8, Fig. 1C) corresponding to collapsed forks^7^. Total DNA synthesis was largely unaltered at early time points (Fig. S1G, 10-20 min), indicating normal replication prior to collapse. Crucially, θ structures almost completely disappeared (Fig. S1H) and collapse approached 100% (Fig. S1I). Thus, we induced highly efficient leading collapse.

Following collapse, σ structures declined (Fig. 1C) and two products emerged. The major product was high molecular weight species that remained in the well (HMw; Fig. 1B lanes 9-10, Fig. 1D) and resembled double-strand break (DSB) repair products^108,109^ (see Fig. 2, below). We also observed the formation of nicked and supercoiled circular monomers (nCMs and scCMs; Fig. 1B, lanes 9-10, Fig. S1J-K) that were far less abundant than the high molecular weight species (Fig. S1I). Because these circular monomers were absent from the control (Fig. 1B, lanes 1-5), it was unlikely they arose from displacement of the LacR barrier. Instead, their existence suggested a mechanism that converts collapsed molecules into full length circular plasmids (see Fig. 4, below). Overall, seDSBs from leading collapse are efficiently converted into at least two products.

**Figure 2:**
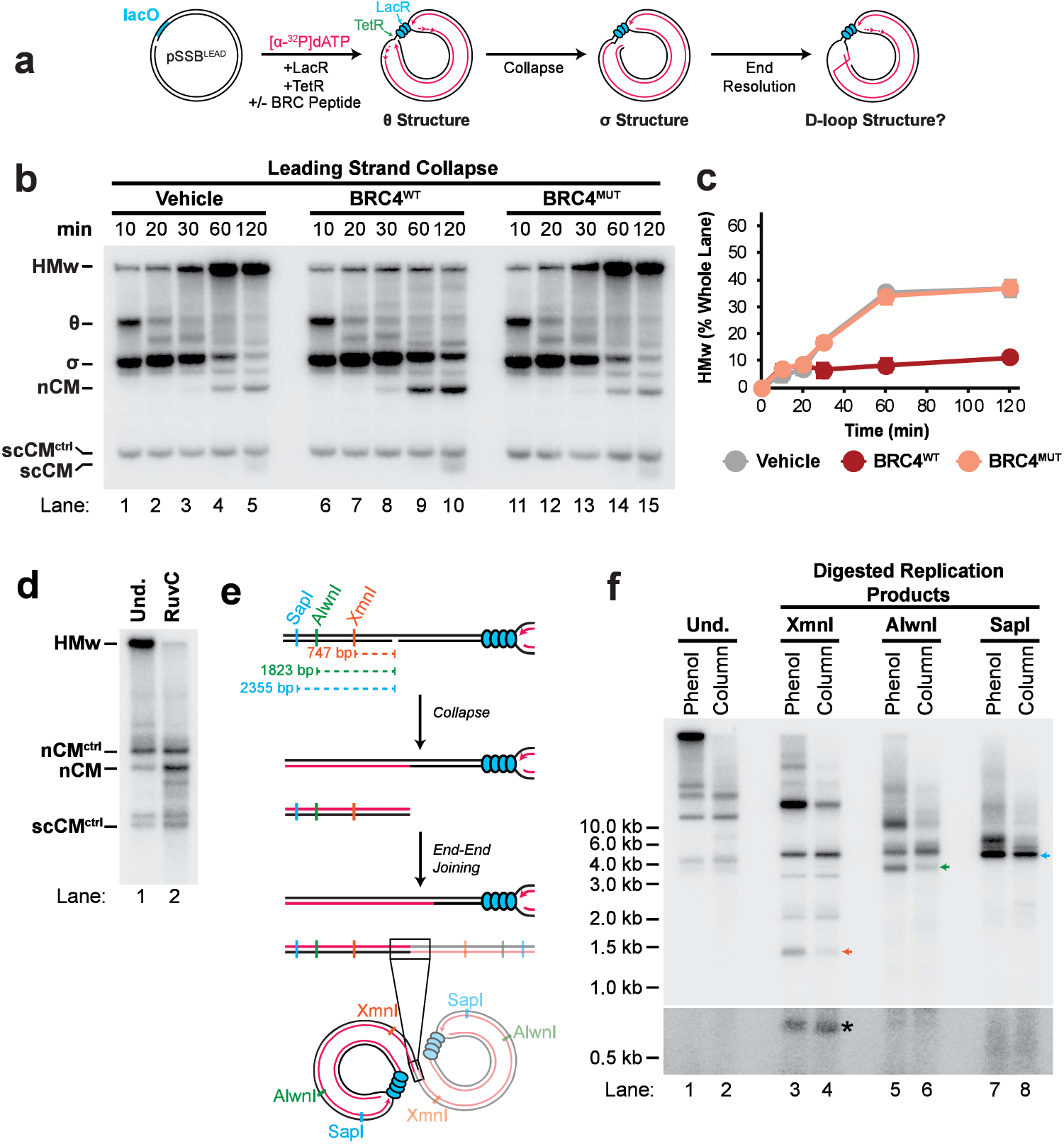
Leading seDSBs undergo RAD51-dependent D-loop formation and end joining. **(A)** Leading collapse was induced by replication of pSSB^LEAD^ in the presence of TetR, LacR and either vehicle, BRC peptide (BRC4^WT^), or Mutant BRC peptide (BRC4^Mut^) as a negative control. **(B)** Samples from (a) were separated on an agarose gel and visualized by autoradiography. **(C)** Quantification of high molecular weight (HMw) products from (b). Mean ± S.D., n = 3 independent experiments. **(D)** Products of leading collapse were purified and digested with RuvC. Digested samples were then separated on an agarose gel and visualized by autoradiography. See also Fig. S2E-H. **(E)** Cartoon depicting expected products of end-to-end fusion events. Digestion of end-to-end fused molecules was expected to generate linear fragments twice the length of the distance between restriction sites and the SSB. **(F)** Products of leading collapse were purified with phenol:chloroform extraction to recover total DNA products or using spin columns to exclude high molecular weight species. Purified DNA samples were then restriction digested, separated on an agarose gel, and visualized by autoradiography. Colored arrows indicate end-to-end linear fragments indicated in (e). Asterisk indicates collapsed arm fragments. See also Fig. S2E-H.

seDSBs can restart replication^6,55–65^. To assess restart efficiency following leading collapse, we used restriction digests to measure nascent DNA synthesis around the leading SSB (collapse region DNA synthesis; Fig. 1E, Fig. S1M). Collapse reduced synthesis ∼4-fold in this region (Fig. 1F,G), matching levels expected if no restart had occurred (Fig. 1G). This was not due to TetR blocking restart because adding tetracycline after collapse did not increase DNA synthesis (Fig. S1N-R). It was also not due to interlinked repair intermediates because the result held true when denaturing alkaline gel electrophoresis was performed (Fig. S1S-U). Notably, this analysis detected the broken end even after two hours (arm; Fig. S1T), which indicated that the seDSB was highly stable. Additionally, our native and denaturing analyses both detected fragment sizes consistent with end-to-end fusions of seDSBs (E-E; Fig. 1F, Fig. S1T; see Fig. 2, below, for further analysis). Thus, replication restart following leading collapse is highly inefficient, despite efficient resolution of seDSBs.

### Collapsed forks form D-loops and undergo end-joining

Following collapse, RAD51 is thought to convert seDSBs to D-loops^55,78^. To test the role of RAD51 in resolving leading seDSBs (Fig. 1), we induced fork collapse in extracts containing a BRCA2-derived peptide that inhibits RAD51 binding to chromatin (BRC4^WT^; Fig. 2B)^103,110,111^ or a mutant peptide that is not thought to disrupt RAD51 function (BRC4^Mut^; Fig. 2B)^103,110^. RAD51 inhibition greatly reduced high molecular weight species (HMw; Fig. 2B, lanes 6-10, Fig. 2C, Fig. S2A) and stabilized collapsed forks (σs; Fig. 2B, lanes 6-10, Fig. S2A), suggesting RAD51 converts collapsed forks to high molecular weight species. Intriguingly, circular monomers increased upon RAD51 inhibition, indicating that RAD51 typically suppresses these products (Fig. 2B, lanes 8-10; Fig. S2B). RAD51 inhibition did not reduce collapse region DNA synthesis (Fig. S2C-D), supporting the conclusion that seDSB resolution does not restart replication. High molecular weight species were resolved by RuvC treatment (HMw; Fig. 2D), indicating they contain 4-way junctions. These 4-way junctions correspond to D-loops, rather than reversed forks or Holliday junctions, because they arise from RAD51-dependent repair of an seDSB. Altogether RAD51 plays two roles following fork collapse: it converts seDSBs to D-loops; and it suppresses formation of full length molecules.

Products of the expected size for end-to-end fusions followed leading collapse (E-E; Fig. 1F) and were suppressed by RAD51 inhibition (E-E; Fig. S2C), suggesting they were RAD51-dependent. To test whether the high molecular weight species contained end-to-end fusions, we used column purification to remove the high molecular weight products, as previously described^112^ (Fig. S2F-G). Restriction digests with enzymes located different distances from the collapse site (Fig. 2E) produced fragments consistent with end-to-end fusions (Fig. 2F, lanes 1,3,5,7) and these were greatly reduced when high molecular weight species were removed (Fig. 2F, lanes 2,4,6,8). These products appeared late (red arrows; Fig. S2G, lanes 4, 8) after the collapsed fork was resolved (collapsed arm; Fig. S2G, lanes 1-3, 5-7). They also could not be resolved by topoisomerase II treatment (Fig. S2H, lane 3), indicating that they did not arise from DNA intertwines. Thus, a fraction of seDSBs are converted to end-to-end fusions by RAD51.

### Lagging seDSBs form D-loops and end-to-end fusions, but do not restart synthesis

We next examined seDSBs that arose from an SSB on the lagging strand template (‘lagging collapse’). We generated pSSB^LAG^ in which the collapse region is the reverse complement of pSSB^LEAD^ (Fig. S1A) so that SSBs were generated on the lagging strand template (Fig. 3A, Supp Fig. 1E). Lagging collapse generated seDSBs (σs; Fig. 3B-C) and collapse efficiency approached 100% (Fig. S3A-B). Lagging seDSBs were converted to high molecular weight species (HMw; Fig. 3B, D, Fig. S3C) containing end-to-end fusions (Fig. S3D-G) and 4-way junctions (Fig. S3H). Crucially, no DNA synthesis was detected in the collapse region (Fig. 3G). Thus, lagging collapse mirrors the consequences of leading collapse (Fig. 1, Fig. 2). However, several unique properties of leading collapse were noted: circular monomers were not detected (Fig. 3B); θs remained detectable, which indicated collapse was less efficient (θs; Fig. 3B, lanes 6-10, Fig. 1B, lanes 6-10); and total DNA signal was lower (Fig. S3I, Fig. S1G), suggesting that more degradation took place. Thus, both leading and lagging collapse generate seDSBs that are resolved to high molecular weight species without detectable replication restart, although some differences exist.

**Figure 3:**
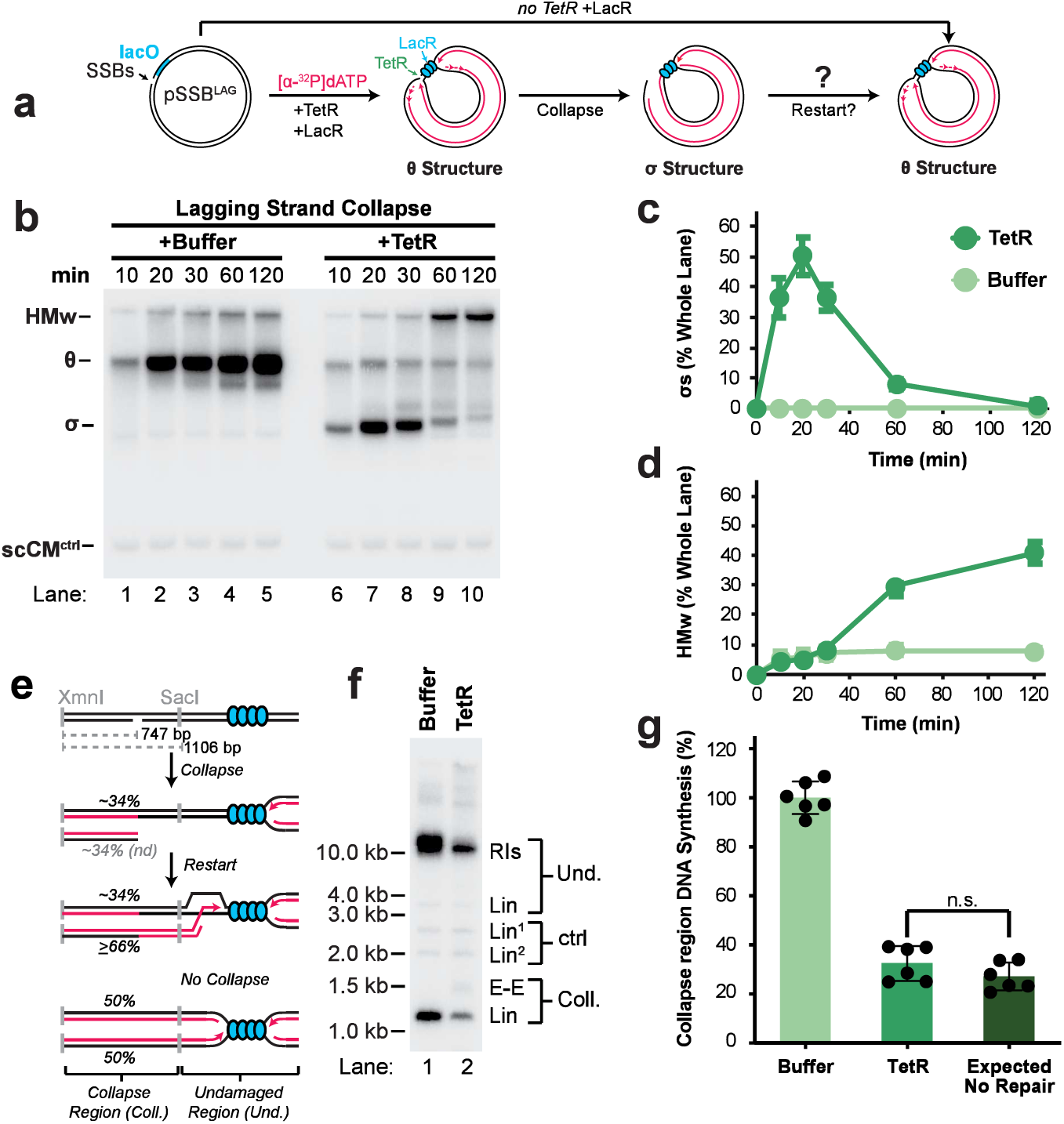
Lagging seDSBs are resolved but do not restart DNA synthesis. **(A)** pSSB^LAG^ was replicated using *Xenopus* egg extract to induce lagging collapse as for leading collapse in Fig. 1A. See also Fig. S1F. **(B)** Samples from (a) were separated on an agarose gel and visualized by autoradiography. **(C)** Quantification of σ structures (σs) from (b). Mean ± S.D., n = 3 independent experiments. **(D)** Quantification of high molecular weight (HMw) products from (b). Mean ± S.D., n = 3 independent experiments. **(E)** Cartoon depicting the assay for restart of DNA synthesis in the collapse region. See also Fig. S1M. **(F)** Purified DNA samples from T=120 in (b) were digested with XmnI and SacI, then separated on an agarose gel and visualized by autoradiography. **(G)** Quantification of collapse region DNA synthesis from (f) as in (e). Signal was normalized to control fragment ‘Lin^1^’. Expected values for no repair account for replication efficiency and collapse efficiency. See also Fig. S1M. Mean ± S.D., n = 6 independent experiments. Data were analyzed by one-way analysis of variance (ANOVA) and Dunnett’s multiple-comparison method.

### Leading SSBs cause secondary replication fork collapse

Leading collapse yielded full length molecules (Fig. 1B) that were suppressed by RAD51 (Fig. 2B) and absent in lagging collapse (Fig. 3B). We hypothesized that these arose from enhanced progression through the LacR barrier, for example by recruitment of a helicase to the D-loop^57,86^. To test this, we performed restriction digests to analyze signal within the *lacO* array (SacI and KpnI; Fig. 4A). However, leading collapse did not induce detectable DNA synthesis within the *lacO* array (*lacO* region; Fig. 4B, lanes 3-4, Fig. 4D). In contrast, restriction digests that directly measured the abundance of full length molecules (AlwNI; Fig. 4A) revealed an ∼3-fold increase in these species (full length product; Fig. 4B, lanes 1-2, Fig. 4C). Hence, leading collapse produced full length molecules without synthesis through the LacR barrier.

**Figure 4:**
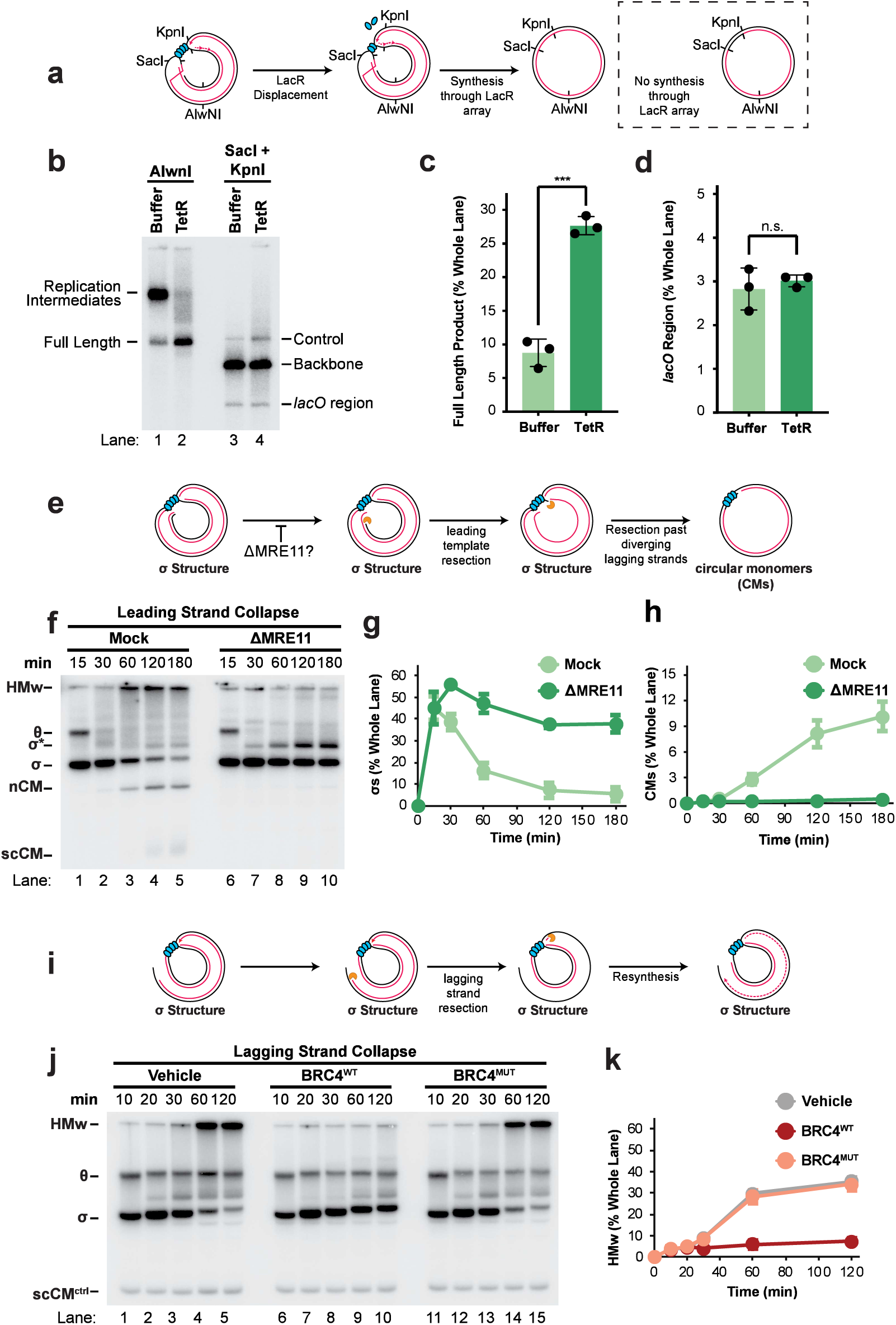
Leading seDSBs can be resolved by extensive nuclease digestion. **(A)** Cartoon depicting formation of full length molecules by synthesis through the LacR array or independent of LacR array displacement. **(B)** Leading collapse was induced as depicted in Fig. 1A. Samples from T=120 min were then purified and analyzed by restriction digest using either AlwNI alone or SacI and KpnI combined. Digested samples were separated on an agarose gel and visualized by autoradiography. **(C)** Quantification of full length linear products that arose from AlwNI digestion in (b). Mean ± S.D., n = 3 independent experiments. Data were analyzed by unpaired t-test. **(D)** Quantification of linearized *lacO* array fragments that arose from SacI-KpnI digestion in (b). Mean ± S.D., n = 3 independent experiments. Data were analyzed by unpaired t-test. **(E)** Cartoon depicts generation of circular monomers (CMs) by extensive degradation of seDSBs. **(F)** Leading collapse was induced as depicted in Fig. 1A in mock- or MRE11-immunodepleted extracts. Samples from were separated on an agarose gel and visualized by autoradiography. **(G)** Quantification of σ structures (σs) from (f). Mean ± S.D., n = 3 independent experiments. **(H)** Quantification of nicked plus supercoiled circular monomers (CMs) from (f). Mean ± S.D., n = 3 independent experiments. **(I)** Cartoon depicts the retention of σ structures (σs) in response to extensive degradation of a lagging seDSB due to resynthesis of lagging strands. **(J)** Lagging collapse was induced as depicted in Fig. 3a. in the presence of either vehicle or BRC peptide. Samples from were separated on an agarose gel and visualized by autoradiography. **(K)** Quantification of high molecular weight products (HMw) from (i). Mean ± S.D., n = 3 independent experiments.

We next hypothesized that nuclease activity was responsible for generation of full length molecules without synthesis through the *lacO* array. In this model, extensive exonuclease activity would remove the *lagging* strand template from the divergent fork, which would cause it to collapse^7^ (Fig. 4E). This ‘secondary collapse’ model predicts that blocking exonuclease activity should block formation of full length molecules (Fig. 4E). We therefore immunodepleted MRE11 to inhibit exonuclease activity at DSBs. MRE11 depletion increased DNA synthesis following collapse (Fig. S4A) and stabilized collapsed forks (σs; Fig. 4F-G; collapsed arm; Fig. S4B-C), demonstrating that exonuclease activity was severely inhibited. MRE11 depletion also blocked formation of high molecular weight species (Fig. 4F, Fig. S4D), consistent with a requirement for exonuclease activity to form D-loops. Importantly, MRE11 depletion also abolished formation of full length molecules (CMs; Fig. 4F, Fig. 4H, Fig. S4E). Thus, extensive exonuclease degradation causes secondary collapse at the divergent fork to generate full length molecules without completion of DNA synthesis.

A further prediction of the ‘secondary collapse’ model is that it should not occur with lagging collapse because this would not affect the template strand of the divergent fork (Fig. 4I). Indeed, no full length molecules formed with lagging collapse (Fig. 3). RAD51 was found to limit secondary collapse at *leading* seDSBs (Fig. 2B), so we inhibited RAD51 with BRC4^WT^ at *lagging* seDSBs to maximize the chances of detecting secondary collapse (Fig. 4J). As with leading collapse (Fig. 2A-C), RAD51 inhibition blocked formation of high molecular weight species (HMw; Fig. 4J-K) and stabilized collapsed forks (σs; Fig. 4J), indicating that D-loop formation was blocked. Collapse region synthesis was unaffected, as expected (Fig. S4F-H). However, no circular monomers were generated following RAD51 inhibition (Fig. 4J, lanes 6-10). Hence, these data establish secondary collapse as a unique mechanism of seDSB resolution that can operate following leading collapse.

### Efficient end resolution and completion of DNA synthesis following bidirectional collapse

Fork convergence at an SSB (‘bidirectional collapse’) can form a double-ended DSB (deDSB) when RAD51 activity is unavailable or an SSB is generated by DNA-protein crosslink^52,77^. However, it is unclear whether this occurs at simple nicks or in the presence of RAD51, when BIR should be available. It is also unclear how effectively the deDSB can complete DNA synthesis. We therefore asked whether fork convergence at an SSB (‘bidirectional collapse’) could trigger deDSB formation and completion of DNA synthesis through double-strand break repair. We replicated pSSB^LEAD^ in the absence of LacR to allow fork convergence at an SSB (pSSB; Fig. 5A). Without TetR, SSBs were quickly repaired, and the expected circular monomeric products of DNA replication formed (nCMs and scCMs; Fig. 5B)^7,107^. With TetR, DNA synthesis increased slightly (Fig. S5A), possibly due to degradation and resynthesis of the parental strand, which would not be radiolabeled in the control. It was not due to re-replication events^27^ because DNA synthesis halted at the same time as the control (Fig. S5A). Importantly, we detected linear molecules that corresponded to formation of a deDSB (Fig. 5B, lanes 6-7, Fig. S5B) and are the expected products of bidirectional collapse (Fig. 5A). Collapse approached 100% (Fig. S5C) and produced high molecular weight species (Fig. 5C), as for leading and lagging collapse (Fig. S1L, Fig. S3C). Thus, bidirectional collapse forms a deDSB that is efficiently resolved.

**Figure 5:**
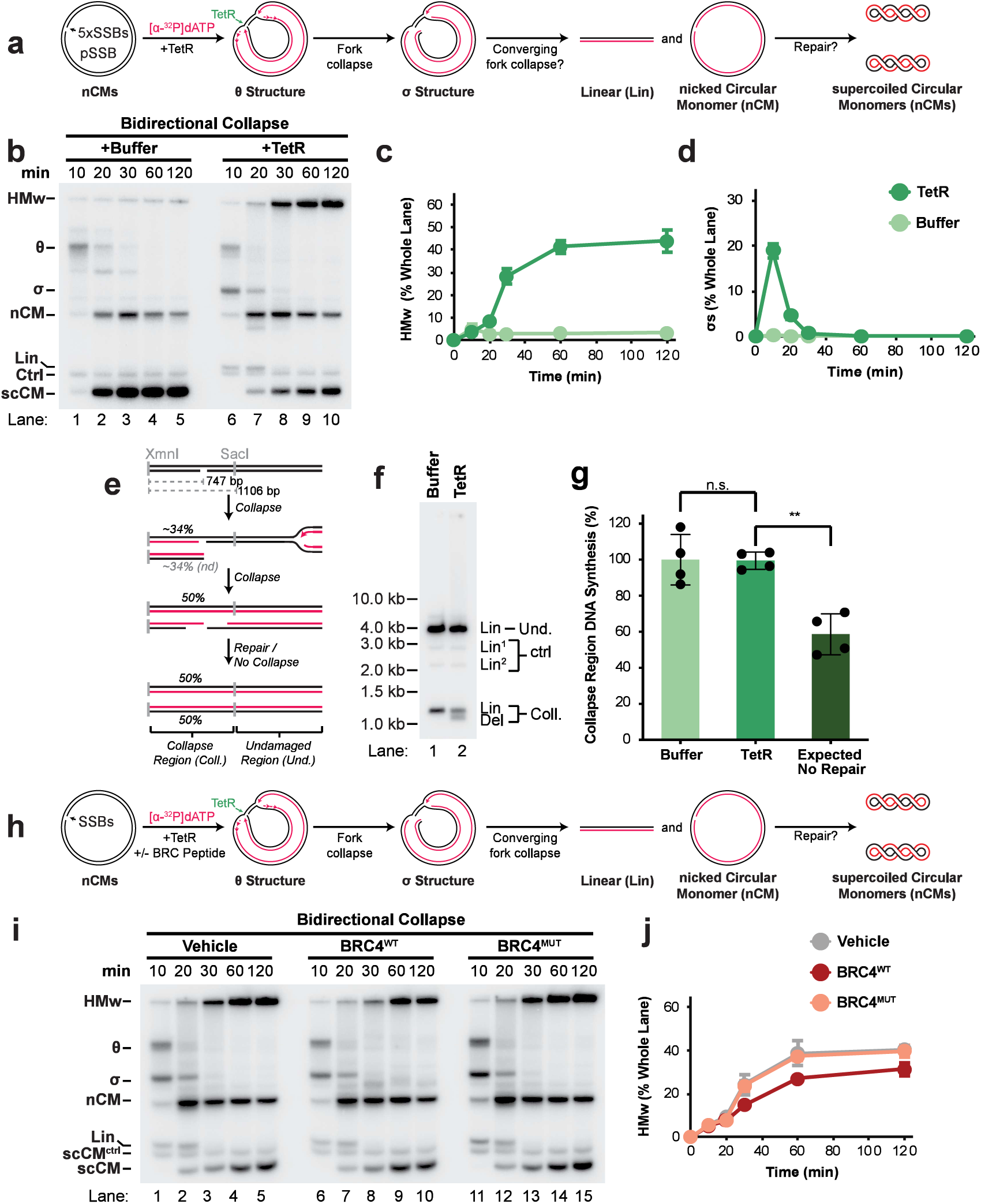
deDSBs arising from fork collapse readily complete DNA synthesis. **(A)** pSSB was replicated using *Xenopus* egg extracts in the presence of (+TetR) to stabilize the SSBs. TetR was omitted in the buffer control (+Buffer), which allowed religation of SSBs prior to replication. A LacR barrier was omitted so that forks could converge upon the SSBs to elicit bidirectional collapse. Nascent strands were radiolabeled by inclusion of [α-^32^P]dATP. Fully replicated plasmid DNA (scCM^ctrl^) served as a loading control. **(B)** Samples from (a) were separated on an agarose gel and visualized by autoradiography. See also Fig. S5A-C. **(C)** Quantification of high molecular weight products (HMw) from (b). Mean ± S.D., n = 4 independent experiments. **(D)** Quantification of σ structures (σs) from (b). Mean ± S.D., n = 4 independent experiments. **(E)** Cartoon depicting the assay for completion of DNA synthesis in the collapse region. **(F)** Purified DNA samples from T=120 in (b) were digested with XmnI and SacI, then separated on an agarose gel and visualized by autoradiography. **(G)** Quantification of collapse region DNA synthesis from (f) as in (e). Signal was normalized to control fragment ‘Lin^1^’. Expected values for no repair account for replication efficiency and collapse efficiency. See also Fig. S1M. Mean ± S.D., n = 4 independent experiments. Data were analyzed by one-way analysis of variance (ANOVA) and Tukey’s multiple-comparison method. **(H)** Bidirectional collapse was induced by replication of pSSB in the presence of TetR as well as either vehicle, BRC peptide (BRC4^WT^), or Mutant BRC peptide (BRC4^Mut^) as a negative control. **(I)** Samples from (h) were separated on an agarose gel and visualized by autoradiography. **(J)** Quantification of high molecular weight products (HMw) from (i). Mean ± S.D., n = 3 independent experiments.

During bidirectional collapse, we also detected seDSBs (σs; Fig. 5B lanes 6-7), which arise from leading or lagging collapse (Fig. 1, Fig. 3, Fig. 5A). seDSBs peaked at 10 minutes and rapidly declined by 20 minutes (σs; Fig. 5B,D) while deDSBs persisted from 10 to 20 minutes, then declined (lins; Fig. 5B, Fig. S5B). Thus, seDSBs were resolved prior to deDSBs. During bidirectional collapse, seDSBs were ∼2.5 fold lower than for leading and lagging collapse (∼20% in Fig. 5D; ∼50% in Fig. 1C, Fig. 3C), suggesting rapid conversion to deDSBs by the converging fork. deDSBs were also 3-fold less abundant than seDSBs (∼6% in Fig. S5B; ∼20% in Fig. 5D) suggesting that deDSBs were more readily resolved than seDSBs. Accordingly, deDSBs arising from bidirectional collapse were resolved by 30 minutes (linears; Fig. S5B) while seDSBs from leading or lagging collapse did not resolve until after 60 minutes (σs; Fig. 1C, Fig. 3C). These data provide direct evidence for the model that bidirectional collapse initially forms an seDSB that the converging fork converts to a deDSB (Fig. 5A) and also suggest that deDSBs resolve more rapidly than seDSBs.

We next determined whether DNA synthesis completed efficiently after bidirectional collapse. We measured DNA synthesis in the collapse region (Fig. 5E) as for leading and lagging collapse (Fig. 1E, Fig. 3E). Surprisingly, collapse region DNA synthesis was indistinguishable from the control (coll; Fig. 5F-G). Furthermore, the high molecular weight products of fork collapse (HMw; Fig. 5B-C) contained only full length molecules (Fig. S5D-F). These data indicated that collapse region synthesis was highly efficient and involved effective daughter molecule separation. Thus, bidirectional collapse enables efficient completion of DNA synthesis through double-strand break repair.

We next examined the repair products of bidirectional collapse. Inspection of the collapse region (coll; Fig. 5F) revealed slightly smaller products (Lin^2^; Fig. 5F) suggesting that small deletions occurred. These deletion products were not detected following leading or lagging collapse (Fig. 1F, Fig. 3F). Similarly, end-to-end fusions arising from leading or lagging collapse (Fig. 1F, Fig. 3F) were not detected for bidirectional collapse (Fig. 5F). Thus, deletions were specific to bidirectional collapse while end-to-end fusions were specific to leading and lagging collapse. We then examined the role of homologous recombination by inhibiting RAD51 activity (Fig. 5H) as for leading and lagging collapse previously (Fig. 2A-C, Fig. 4I-K). Formation of high molecular weight species was reduced but not blocked (Fig.5I, lanes 6-10, Fig. 5J), suggesting a limited requirement for homologous recombination. Accordingly, these high molecular weight products were only partially sensitive to RuvC (Fig. S5G), suggesting limited D-loop formation. Because these species were fully resolved by restriction digest but not topoisomerase treatment (Fig. S5H) they were likely concatemers formed by end joining. RAD51 inhibition did not affect collapse region DNA synthesis (Fig. S5I-K) or deletion events (Del; Fig. S5J). Thus, bidirectional collapse can lead to efficient completion of DNA synthesis independent of homologous recombination, likely by end joining. These data are consistent with the ability of both homologous recombination and end-joining pathways to act at deDSBs^52,79^, but differ from the prevailing view that DSB resolution at collapsed forks is critically dependent on homologous recombination^4,6,9,42,51–55,78^.

### End resolution is separate from completion of DNA synthesis following Cas9 induced collapse

We next tested whether our findings from ‘simple’ SSBs apply to CRISPR-Cas9 SSBs, which other recent studies have examined^51,52^. We employed the H840A Cas9 nickase (nCas9), because it primarily generates seDSBs^52^, as we observed for leading and lagging collapse at simple SSBs (Fig. 1, Fig. 3). We avoided D10A Cas9 because it generates deDSBs following lagging collapse and this would complicate our analysis^52^. We constructed a plasmid harboring multiple nCas9 target sites adjacent to a *lacO* array, akin to the design of pSSB^LEAD^ and pSSB^LAG^ (Fig. 1, Fig. 3). We replicated this plasmid (pCRISPR; Fig. 6A) with either nCas9 or nuclease-dead mutant (dCas9), plus LacR and the appropriate guide RNA to target the leading strand template (Fig. 6A). We also omitted LacR so we could directly compare leading and bidirectional collapse (Fig. 6A).

**Figure 6:**
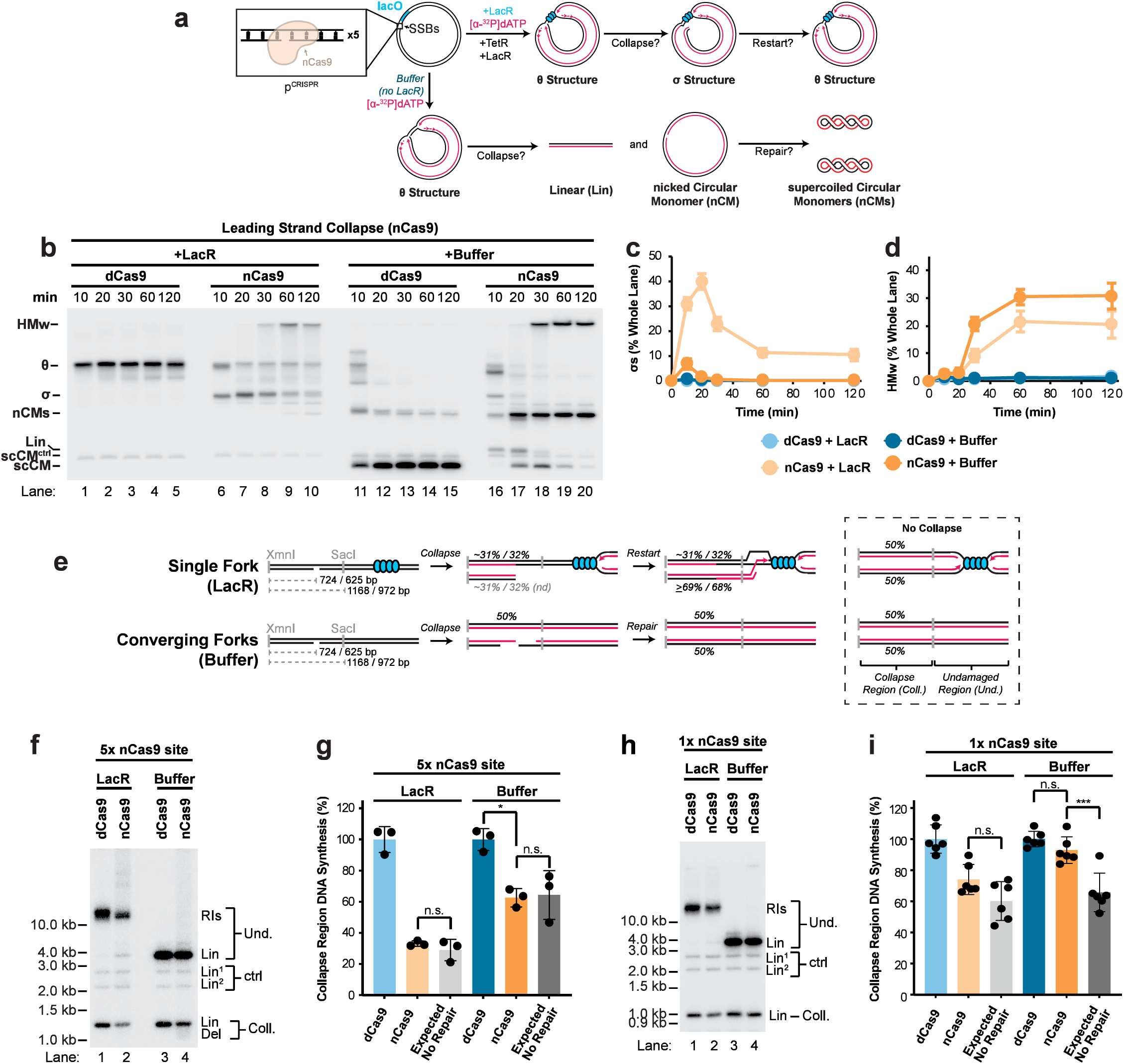
Analysis of nCas9 induced fork collapse. **(A)** pCRISPR was replicated using *Xenopus* egg extracts in the presence of Cas9^D840A^ nickase (nCas9) to generate SSBs. Nuclease-dead Cas9 (dCas9) did not generate SSBs and was used as a control. LacR was included to induce leading collapse or omitted to induce bidirectional collapse. Nascent strands were radiolabeled by inclusion of [α-^32^P] dATP. Fully replicated plasmid DNA (scCM^ctrl^) served as a loading control. **(B)** Samples from (a) were separated on an agarose gel and visualized by autoradiography. **(C)** Quantification of σ structures (σs) in (b). Mean ± S.D., n = 3 independent experiments. **(D)** Quantification of high molecular weight products (HMw) in (b). Mean ± S.D., n = 3 independent experiments. **(E)** Cartoon depicting the assay for restart/completion of DNA synthesis in the collapse region. Numbers indicate the expected DNA signal for pCRISPR (used above) followed by pCRISPR^1X^ (used below). **(F)** Purified DNA samples from T=120 in (b) were digested with XmnI and SacI, then separated on an agarose gel and visualized by autoradiography. See also Fig. S6T-V. **(G)** Quantification of collapse region DNA synthesis from (f) as in (e). Signal was normalized to control fragment ‘Lin^1^’. Expected values for no repair account for replication efficiency and collapse efficiency. Mean ± S.D., n = 3 independent experiments. Data were analyzed by one-way analysis of variance (ANOVA) and Tukey’s multiple-comparison method. **(H)** Fork collapse was induced as in (a) using pCRISPR^1X^ that contained only a single copy of the nCas9 target sequence. Purified DNA samples from T=120 in Fig. S6T were digested with XmnI and SacI, then separated on an agarose gel and visualized by autoradiography. **(I)** Quantification of collapse region DNA synthesis from (h) as in (e). Signal was normalized to control fragment ‘Lin^1^’. Expected values for no repair account for replication efficiency and collapse efficiency. Mean ± S.D., n = 6 independent experiments. Data were analyzed by one-way analysis of variance (ANOVA) and Tukey’s multiple-comparison method.

nCas9 induction of leading or bidirectional collapse generated seDSBs (σs; Fig. 6B, lanes 6-10, 16-20, and Fig. 6C), as expected (Fig. 1, Fig. 5). Collapse was also efficient in both cases (∼80%; Fig. S6A-E), but lower than for TetR-stabilized SSBs (∼100%; Fig. S1I, Fig. S5C). Leading collapse produced seDSBs that were converted to high molecular weight species (Fig. 6B, lanes 6-10, Fig. 6C-D) as expected (Fig. 1, Fig. 3). Bidirectional collapse produced fewer seDSBs that were converted to deDSBs and then high molecular weight species (σs, Lins, HMws; Fig. 6B, lanes 16-20, Fig. 6C-D), also as expected (Fig. 5). High molecular weight species from leading collapse were sensitive to RAD51 inhibition(Fig. S6F-J) and RuvC treatment (Fig. S6P-Q, lanes 1-2). High molecular weight species from bidirectional collapse were less affected by RAD51 inhibition (Fig. S6K-O) and RuvC treatment (Fig. S6P, lanes 3-4) as expected (Fig. 2A-C, Fig. 5H-J). Full length products were generated following leading collapse (Fig. 6B, lanes 6-10) but not in the nuclease dead control (Fig. 6B, lanes 1-5, Fig. S6R) or following nCas9 induced lagging collapse (Fig. S6S), indicating that secondary collapse occurred following leading collapse induced by nCas9. Hence, the DNA structures generated by nCas9 collapse closely resembled those formed at simple SSBs (Figs. 1-5).

We next monitored restart and completion of DNA synthesis following nCas9-induced collapse (Fig. 6E). Leading collapse did not result in detectable synthesis in the collapse region (Fig. 6F, lanes 1-2, Fig. 6G) as expected (Fig. 1E-G). However, we did not detect the end-to-end fusions (Fig. 6F) that were observed at simple SSBs (Fig. 1F). Surprisingly, nCas9-induced bidirectional collapse also did not result in detectable synthesis in the collapse region (Fig. 6F lanes 3-4, Fig. 6G) in contrast to bidirectional collapse at simple SSBs (Fig. 5E-G). Yet, DSB resolution was highly efficient for nCas9-induced leading and bidirectional collapse (Fig. 6A-D). These results strongly support the notion that DSB resolution and completion of DNA synthesis are separate events.

Although collapse region DNA synthesis was not detected for nCas9-induced bidirectional collapse (Fig. 6F-G), we detected a very low level of deletion products (Del; Fig. 6F lane 4), implying a low level of repair. Because Cas9 remains associated with DSBs^113^ and inhibits their repair^114^, the nCas9 target sites in pCRISPR could have interfered with completion of DNA synthesis (Fig. 6F-G). In support of this idea, seDSBs persisted longer when induced by nCas9 (Fig. 6C) than in response to simple SSBs (Fig. 1C). We therefore induced leading and bidirectional collapse at a single nCas9 target site (Fig. 6H-I). Under these conditions, leading collapse and bidirectional collapse were inefficient but detectable (Fig. S6T-W). Collapse region DNA synthesis was detected for bidirectional collapse (Buffer; Fig. 6I) but not leading collapse (LacR; Fig. 6I). These experiments were highly variable (Fig. 6I), suggesting complex and stochastic repair events. Nonetheless, these results support the conclusion that bidirectional collapse efficiently completes DNA synthesis while leading collapse does not, as for simple SSBs (Fig. 1E-G, Fig. 3E-G).

### Fork collapse following PARP inhibition is flexible

Abasic sites (Apurinic/Apyrimidinic sites; APs) are a common form of DNA damage that can be converted to ‘complex’ SSBs. Fork collapse at AP sites enhances the vulnerability of cancer cells to PARP inhibition^21,22^. To study AP site replication in *Xenopus* egg extracts we previously generated a plasmid containing an AP site (pAP) that was flanked by *lacO-*bound LacR to prevent repair prior to replication^115^. However, when this AP plasmid was introduced into extracts, it led to formation of a durable SSB (Fig. S7A-B)^115^. We therefore used this ‘AP-SSB’ to investigate fork collapse at a ‘complex’ SSB and the effects of PARPi.

pAP contained a *lacO* array and was replicated in the presence of LacR to ensure unidirectional collapse, in this case lagging collapse, while also stabilizing the AP (Fig. 7A). AP-SSB replication produced prominent seDSBs (σs; Fig. 7B lanes 1-3) that became high molecular weight species (HMw; Fig. 7B lanes 4-5) in a RAD51-dependent manner (Fig. 7B, lanes 6-10, Fig. 7C, Fig. S7C). We then assessed whether the PARP inhibitor talazoparib (PARPi; Fig. S7D) could stabilize AP-SSBs. PARPi did not affect control plasmid replication (Fig. S7E-G) and minimally impacted the formation of AP-SSB-induced seDSBs (Fig. 7D-H). PARPi is effective in extracts^77^ so this was not due to extract-specific effects of PARPi but was unexpected because PARPi should potently induce SSBs at AP sites^21,22,50^. Hence, collapse at AP-SSBs was similar to ‘simple’ SSBs and unexpectedly independent of PARPi.

**Figure 7:**
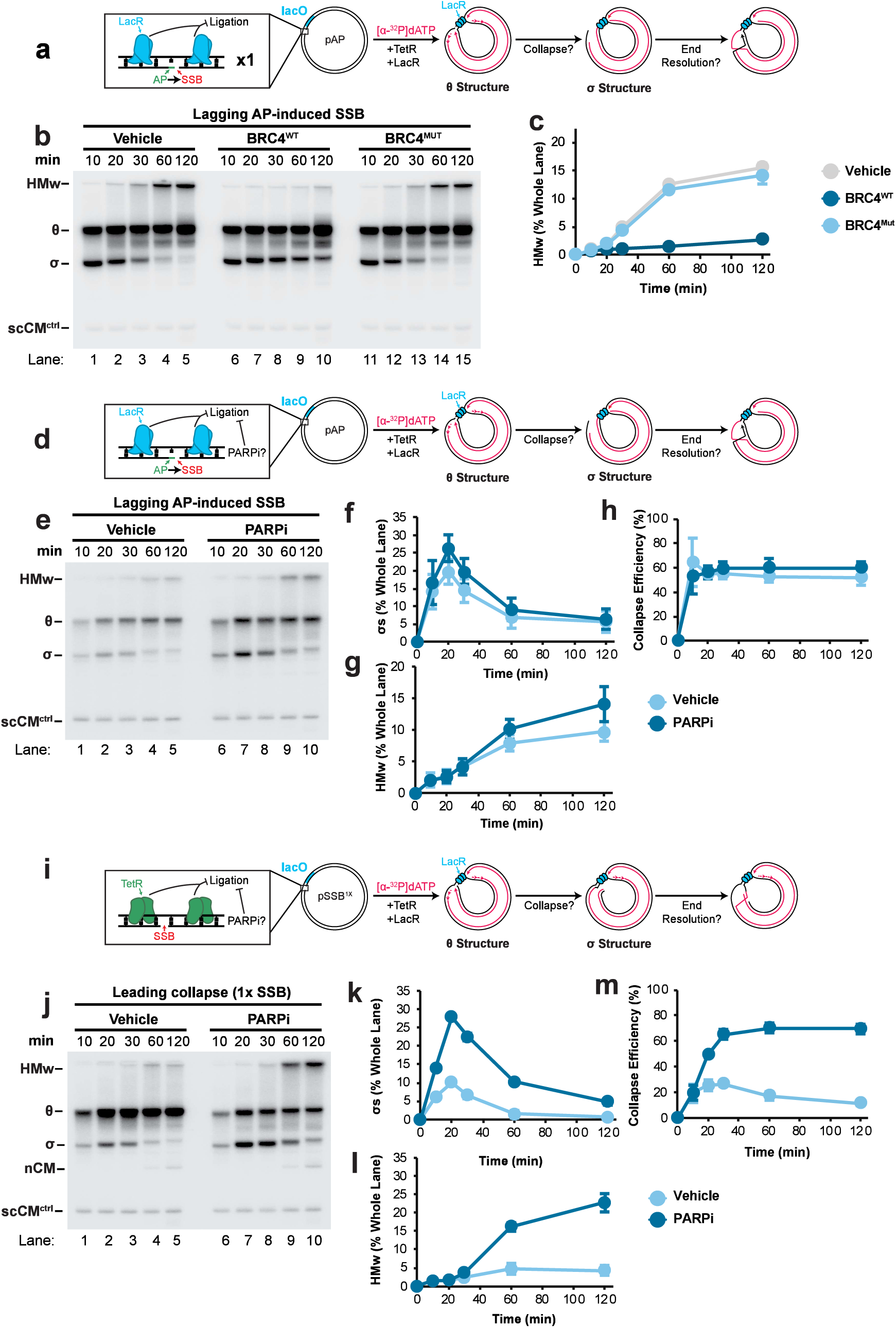
SSB composition influences PARP inhibition. **(A)** pAP, which contains a site-specific abasic site (AP site) on the lagging strand template, was replicated using *Xenopus* egg extract. LacR was included in the reaction to ensure lagging collapse. Reactions were supplemented with either vehicle control, BRC peptide (BRC4^WT^), or Mutant BRC peptide (BRC4^Mut^) as an additional negative control. Nascent strands were radiolabeled by inclusion of [α-^32^P] dATP. Fully replicated plasmid DNA (scCM^ctrl^) served as a loading control. **(B)** Samples from (a) were separated on an agarose gel and visualized by autoradiography. **(C)** Quantification of high molecular weight products (HMw) from (b). Mean ± S.D., n = 3 independent experiments. **(D)** Lagging collapse was induced at pAP, as in (a), in the presence or absence of PARPi. **(E)** Samples from (d) were separated on an agarose gel and visualized by autoradiography. **(F)** Quantification of σ structures (σs) from (e). Mean ± S.D., n = 3 independent experiments. **(G)** Quantification of high molecular weight products (HMw) from (e). Mean ± S.D., n = 3 independent experiments. **(H)** Quantification of collapse efficiency from (e). Mean ± S.D., n = 3 independent experiments. Collapse efficiency was calculated as described in methods. **(I)** pSSB^1X^, which contains a single SSB flanked by *tetO* sites, was replicated in the presence of TetR and LacR to ensure leading collapse. Reactions were supplemented with either vehicle or PARPi. **(J)** Samples from (i) were separated on an agarose gel and visualized by autoradiography. **(K)** Quantification of σ structures (σs) from (j). Mean ± S.D., n = 3 independent experiments. **(L)** Quantification of high molecular weight products (HMw) from (j). Mean ± S.D., n = 3 independent experiments. **(M)** Quantification of collapse efficiency from (j). Mean ± S.D., n = 3 independent experiments. Collapse efficiency was calculated as described in methods.

The unexpected resistance of AP-SSBs to PARPi led us to test whether simple SSBs could be affected by PARPi. Collapse using pSSB^LEAD^ resulted in approximately 100% collapse, which prevented detection of further increases in collapse. We therefore tested the effect of PARPi on fork collapse at a single SSB by removing four of the five nicking sites on pSSB^LEAD^ to generate pSSB^1X^ (Fig. 7I). Without TetR there was no detectable collapse (Fig. S7H-J). PARPi also had no effect, which was unexpected because the unprotected SSB should be readily detected by PARP^1,39–41^ (Fig. S7 H-J). With TetR, the SSB was stabilized and fork collapse led to modest levels of seDSBs and high molecular weight species (Fig. 7J-M). Strikingly, PARPi synergized with the TetR-stabilized SSB to increase seDSBs and high molecular weight species (Fig. 7J-L), amounting to an approximately 3-fold increase in collapse events (Fig. 7M). seDSB resolution involved RAD51 (Fig. S7J-N), as expected (Fig. 2A-C, Fig. 4I-K). Additionally, nCMs were detected both in the presence and absence of PARPi (Fig. 7J), indicating that secondary collapse occurred in both conditions. Overall, these results show that PARP activity can be crucial to repair a ‘simple’ SSB when flanked by a DNA-binding protein, demonstrating significant flexibility in how PARPi affects fork collapse. Moreover, the occurrence of secondary collapse following PARPi treatment suggests this mechanism may be relevant to cancer therapy.

## Discussion

### Consequences of fork collapse

We interrogated replication fork collapse using both simple single-strand DNA breaks (SSBs) and Cas9 nickase-generated SSBs (Fig. 8Ai-iv). Collapse from both types of SSBs was similar (Figs 1-3, Fig. 5) and collapse at simple SSBs was enhanced by PARP inhibitor treatment (Fig. 7I-M), which strengthens the relevance of our study to cancer therapy. Following collapse, we found that double-strand breaks (DSBs) produced by fork collapse were efficiently resolved, regardless of whether the SSB was on the leading or lagging strand template, or between converging forks (Fig. 8B-D). Importantly, we provide direct evidence that fork convergence at an SSB involves formation of a single-ended DSB (seDSB) that is converted to a double-ended DSB (deDSB; Fig. 5A-B), consistent with previous results^51–53,56^.

**Figure 8:**
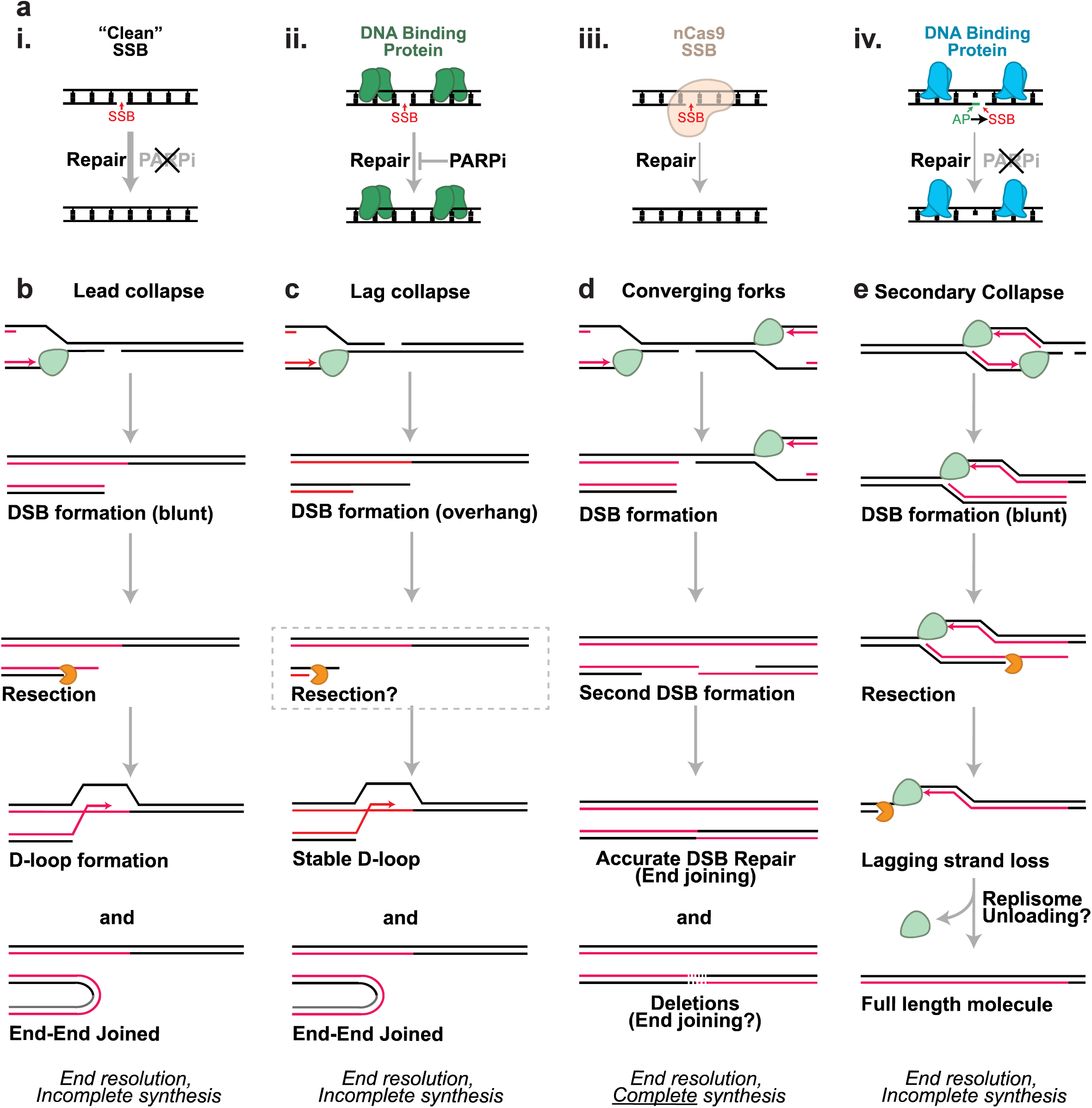
Models for SSB formation and replication fork collapse in *Xenopus* egg extracts. (a) ‘Simple’ SSBs arising from a discontinuity in the phosphodiester backbone are readily repaired in a PARPi-resistant manner (i). However, adjacent DNA-binding proteins can slow repair and confer sensitivity of repair to PARPi in the presence of DNA-binding proteins (ii). nCas9 generated SSBs are repaired slowly (iii). AP site induced SSBs are also repaired slowly and this can be resistant to PARPi (iv). Arrow thickness indicates repair speed. (b) Leading collapse leads to formation of a blunt DSB that is resected and then converted to a stable D-loop. End-to-end joined products are an alternative, erroneous outcome. (c) Lagging collapse proceeds similarly to leading collapse, except that a DNA overhang is initially formed and the requirement for resection is untested. (d) After a fork collapses at an SSB, arrival of the opposing fork can generate a second DSB. The resulting two-ended DSB can be efficiently repaired, likely through end joining. Deletion products are an alternative, erroneous outcome. (e) Following leading collapse, the DSB can undergo extensive resection that removes the broken arm. This could occur if resection proceeded beyond the opposing fork to remove the lagging strand template, which would be expected to result in replisome unloading. Arrival of forks from adjacent origins could then allow for error free replication of the collapse region (not shown).

Resolution of leading and lagging collapse primarily occurred through formation of stable D-loops without detectable restart of DNA synthesis (Fig. 8B-C). By contrast, bidirectional collapse led to efficient completion of DNA synthesis via double strand break repair (Fig. 8D). Hence, DSBs from leading and lagging collapsed forks were resolved without apparent replication restart, whereas DSBs from bidirectional collapse forks were resolved and completed DNA synthesis. These findings reveal a mechanistic separation of end resolution from completion of DNA synthesis following fork collapse. This distinction is further supported by our discovery of secondary collapse, which resolves DSBs without completing DNA synthesis (Fig. 8E, see below). Thus, resolution of collapsed forks is separate from completion of DNA synthesis.

Our observations are highly complementary to those of Winterhalter et al,^116^ who examined fork collapse in bacteria. Both our study and theirs found that fork collapse engages the homologous recombination machinery to form a D-loop, which cannot efficiently restart DNA synthesis. In Winterhalter et al, this is because replicative helicase re-loading is required, while in our study it is because fork convergence at the SSB is necessary. These differences make intuitive sense: the bacterial helicase reloading proteins^117^ are not conserved in vertebrates, and converging forks are rare in bacteria but common in vertebrates^118^. Thus, the events that follow fork collapse may have been shaped by evolution.

### Fate of DSBs arising from fork collapse

seDSBs formed RAD51-dependent D-loops that persisted for hours (Fig. 1, Fig. 3). In contrast, deDSBs were resolved largely independently of RAD51 (Fig. 5). These findings suggest that seDSBs were stabilized by homologous recombination while deDSBs were rapidly repaired by end joining. Although most previous studies did not implicate end joining in resolution of deDSBs following fork collapse (^4,6,9,42,51–55^ but also ^56,57,80–84^), our results suggest that deDSBs arising from fork collapse can undergo end-joining, as for replication-independent deDSBs^52,79^. Two explanations may account for why this was not detected previously. First, several studies were performed in yeasts that inherently favor homologous recombination^6,51,56^. Second, arrival of the converging fork was delayed in mammalian cells^52^, compared to our system (Fig. 5). The longer duration between initial seDSB formation and fork convergence may have led to extensive degradation of the seDSB, which might preclude end joining. It will be important to determine how the timing of fork convergence influences which DSB repair pathways operate following fork collapse.

Our inability to detect break-induced replication (BIR) is consistent with evidence that most fork collapse events involve deDSB formation^51–53,75,76^. Additionally, PARP inhibitors induce fork collapse^11,32–35^ but BIR proteins^67,96^ do not appear to be major determinants of PARP inhibitor cytotoxicity^119^. Thus, seDSBs may not require BIR to complete DNA synthesis. We speculate that BIR is more critical when fork convergence is not possible, as for mitotic DNA synthesis (MiDAS) and alternative lengthening of telomeres (ALT)^66–70^. Nonetheless, BIR is well established^6,55–65^, so its absence in our study suggests something in our system acts as a barrier to BIR. Several considerations lead us to favor the idea that chromatin impedes migration of the D-loop formed by the seDSB in our system. First, no chromatin remodelers would be expected at the D-loop according to known BIR proteins in vertebrates^6,67,96^. Second, a chromatin remodeler was implicated in the response to seDSBs in yeast^51^. Third, chromatin constrains DNA replication^120^ so a chromatin barrier could explain why fork convergence promotes completion of DNA synthesis: the converging fork may displace nucleosomes to allow repair pathways to operate. Another possibility is that BIR operates when single-stranded DNA template is abundant, for example due to R-loops^16,96^ or C-circles^121^. It will be crucial to determine how BIR is deployed.

A notable outcome of leading and lagging collapse is the formation of RAD51-dependent end-to-end fusions, as reported previously^57^. Whether RAD51 acts directly or indirectly is unclear. The *tetO* sequences flanking each collapse site (Fig. 1A) are imperfect palindromes that might allow RAD51-mediated strand annealing^65^. End-to-end fusions were present within the same molecules that contained D-loops (Fig. 2D-F) so they could also be an indirect consequence of D-loop formation. For example, D-loop disruption by negative regulators of RAD51 filaments^122^ may facilitate end joining at seDSBs. However, we disfavor the latter model because an seDSB displaced from a D-loop should contain a single-stranded DNA overhang that should prevent end joining pathways from operating.

Our data reveal that leading collapse can result in complete removal of the seDSB via exonuclease activity to generate a full length molecule (Fig. 8E). We propose a “secondary collapse” model whereby 5’-3’ resection removes the lagging strand template of the diverging fork. Loss of the lagging strand template would trigger replisome removal by the same mechanism that operates during replication termination^7,73^. Secondary collapse would disassemble the replication bubble (Fig. 8E) and allow converging forks from the adjacent origins to replicate the region in a potentially error free manner. Because this mechanism competes with homologous recombination (Fig. 2B) and operates in the presence of PARP inhibitor (Fig. 7I-M), it could feasibly contribute to PARP inhibitor resistance in BRCA-deficient cells.

### Variable effects of PARPi at SSBs

We found that PARP inhibition promoted fork collapse when TetR was bound on either side of a simple SSB but not when the SSB was unbound. Conversely, collapse at AP sites was not enhanced by PARP inhibitors. Both results contradict expectations: it should be possible for simple SSBs to be readily religated, while PARP inhibition should potently induce SSBs at AP sites^39–41,46–50^. Although the generality of these findings is unclear, they demonstrate that fork collapse can be strongly influenced by proteins binding near the SSB. It is thus possible that transcription factor binding in certain cell types (e.g. cancer cells) enhances SSBs and fork collapse. Testing this idea could reveal a new vulnerability to PARP inhibitors.

### Limitations of the study

Our data do not address whether BIR can occur, only that it is not a major outcome in this system. This is because our bulk assays would not detect a minority of BIR events. We cannot discern whether end-to-end fusions at collapsed forks reflect the method of SSB induction (TetR-stabilized simple SSB or CRISPR-Cas9-bound SSB) because each approach is linked to particular DNA sequence. Our results also do not address under which conditions DNA-binding proteins affect resolution of AP sites and SSB repair, only that DNA-binding proteins can regulate SSB repair. Lastly, we cannot exclude the possibility that inefficient restart after an seDSB is unique to the *Xenopus* egg extract system. For instance, these extracts could lack a key BIR protein or proximity to the LacR barrier might prevent DNA synthesis at a D-loop. However, we note that all vertebrate BIR proteins ^57,96,97^ are detectable on DNA in *Xenopus* egg extracts^123^, and replication-coupled repair can readily occur adjacent to a LacR barrier^115,124,125^.

## Supplementary figure legends

<u>**Figure S1: Analyses of leading collapse in *Xenopus* egg extracts**</u>

**(A)** Encounter with an SSB on the leading strand template, results in the formation of a blunt seDSB (leading collapse). **(B)** Encounter with an SSB on the lagging strand template, results in formation of an seDSB with an overhang (lagging collapse). **(C)** Encounter with a lagging SSB can also result in continued unwinding and formation of a deDSB. **(D)** Fork convergence at an SSB can also result in formation of a deDSB. **(E)** To induce leading collapse, pSSB^LEAD^ contained 5x tandem Nb.BsmI nicking sites (‘N1’ to ‘N5’) each flanked by 1x *tetracycline* operator (*tetO*) sequence either side. To induce lagging collapse, pSSB^LAG^ contains the reverse complement of the *tetO*/SSB array from pSSB^LEAD^. **(F)** Cartoon depicting the DNA structures from Fig. 1A along with the loading control, which was replicated prior to replication of pSSB^LEAD^. **(G)** Quantification of total DNA synthesis from Fig. 1B normalized to the maximum signal across all time points and conditions. Mean ± S.D., n = 9 independent experiments. **(H)** Quantification of θ structures from Fig. 1B. Mean ± S.D., n = 9 independent experiments. **(I)** Quantification of collapse efficiency from Fig. 1B. Mean ± S.D., n = 9 independent experiments. Collapse efficiency was calculated as described in methods. **(J)** Quantification of nicked plus supercoiled circular monomers (CMs) from Fig. 1B. Mean ± S.D., n = 9 independent experiments. **(K)** Overexposure of the bottom portion of Fig. 1B to show that scCMs form following collapse (lane 10) but not without collapse (lane 5). **(L)** Quantification of σ structures (σs), high molecular weight products (HMw), and circular monomers (CMs) from Fig. 1B. Mean ± S.D., n = 9 independent experiments. **(M)** Detailed schematic of the expected products arising from the XmnI-SacI digestion depicted in Fig. 1E. Control digestion products arise from digestion of the pre-replicated control plasmid. Fragment lengths are indicated in base pairs (bp) along with the expected radioactive signal relative to the ‘no collapse’ control. **(N)** To address whether TetR binding interferes with seDSB repair, pSSB^LEAD^ was replicated as in Fig. 1B, lanes 6-10. Tetracycline (TC) was added to reactions at the onset of replication (TC-0), at 60 minutes (TC-60) and compared to a no addition control (NA). After replication, samples were separated on an agarose gel and analyzed by autoradiography. TC addition at the onset of replication completely blocked formation of σs (lanes 1, 3, 5), which indicated that TC addition efficiently displaced TetR. **(O)** Quantification of θ structures from (n). Mean ± S.D., n = 3 independent experiments. **(P)** Quantification of σ structures (σs) from (n). Mean ± S.D., n = 3 independent experiments. Note that TC addition at 60 minutes (TC-60) did not impact σ structure resolution, which indicated that residual TetR binding did not impact seDSB resolution. **(Q)** Purified DNA samples from T=120 in (n) were analyzed as in Fig. 1F. **(R)** DNA synthesis in the collapse region was quantified from (q) as in Fig. 1G. Mean ± S.D., n = 3 independent experiments. Data were analyzed by one-way analysis of variance (ANOVA) and Dunnett’s multiple-comparison method. Addition of TC at 60 min did not impact DNA synthesis in the collapse region, which indicated that residual TetR binding did not interfere with seDSB repair. **(S)** To determine whether formation of complex intermediates prevented detection of DNA synthesis in the collapse region in Fig. 1E-G individual DNA strands were analyzed. DNA samples were generated as in Fig. 1F. Graph depicts quantification of total DNA normalized to the maximum signal across all time points and conditions. Mean ± S.D., n = 4 independent experiments. **(T)** Samples from (s) then separated on an alkaline denaturing agarose gel to separate all DNA strands. DNA fragments were then visualized by autoradiography. The broken arm was readily visible, which supported the notion that DNA synthesis did not restart at the seDSB. **(U)** DNA synthesis in the collapsed region was quantified from (t) as in Fig. 1G. Control fragments could not be resolved in (s), so values were normalized to the control fragment ‘Lin^1^’ from the corresponding native gels. Mean ± S.D., n = 4 independent experiments. Data were analyzed by one-way analysis of variance (ANOVA) and Dunnett’s multiple-comparison method. Alkaline denaturing analysis did not detect any synthesis in the collapse region, which indicated that formation of complex intermediates could not explain the absence of detectable DNA synthesis in the collapse region. Note that expected synthesis for no repair was lower than previously due to reduced overall DNA synthesis in this set of experiments, as shown in (s).

<u>**Figure S2: Analyses of RAD51 function during leading collapse**</u>

**(A)** Quantification of DNA synthesis from Fig. 2B normalized to the maximum signal across all time points and conditions. Mean ± S.D., n = 3 independent experiments. **(B)** Quantification of nicked plus supercoiled circular monomers (CMs) from (Fig. 2b). Mean ± S.D., n = 3 independent experiments. **(C)** Purified DNA products from T=120 in Fig. 2B were digested with XmnI and SacI to analyze DNA synthesis in the collapse region. Digested samples were separated on an agarose gel and visualized by autoradiography. **(D)** Quantification of DNA synthesis in the collapse region from (C) normalized to control fragment ‘Lin^1^’. Mean ± S.D., n = 3 independent experiments. Data were analyzed by one-way analysis of variance (ANOVA) and Dunnett’s multiple-comparison method. **(E)** pSSB was replicated in *Xenopus* egg extracts to induce leading collapse as in Fig. 1A. **(F)** Samples from (e) were purified by either phenol:chloroform extraction, to recover total DNA, or spin column purification to exclude high molecular weight DNA. Purified DNA samples were separated on an agarose gel and visualized by autoradiography. High molecular weight species were not detected after column extraction but all other species could be detected, which confirmed that column purification effectively excluded high molecular weight species **(G)** Samples from (f) were digested with AlwNI, then separated on an agarose gel, and visualized by autoradiography. Red arrows indicate fragments enriched in phenol:chloroform purified samples. **(H)** Phenol:chloroform samples from (f) were treated with either AlwNI or Topo II enzyme then separated on an agarose gel and visualized by autoradiography. High Molecular Weight (HMw) species were resolved by AlwNI treatment but not topo II treatment, which indicated that HMw species did not correspond to catenanes.

<u>**Figure S3: Analyses of lagging collapse in *Xenopus* egg extracts**</u>

**(A)** Quantification of θ structures from Fig. 3B. Mean ± S.D., n = 3 independent experiments. **(B)** Quantification of collapse efficiency, as described in the methods, from Fig. 3B. Mean ± S.D., n = 3 independent experiments. Collapse efficiency was calculated as described in methods. **(C)** Quantification of σ structures (σs) and high molecular weight products (HMw) from Fig. 3B. Mean ± S.D., n = 3 independent experiments. **(D)** pSSB^LAG^ was replicated in *Xenopus* egg extracts to induce lagging collapse as in Fig. 3A. **(E)** Samples from (D) were purified by either phenol:chloroform extraction, to recover total DNA, or spin column purification to exclude high molecular weight DNA. Purified DNA samples were separated on an agarose gel and visualized by autoradiography. High molecular weight species were not detected after column extraction but all other species could be detected, which confirmed that column purification effectively excluded high molecular weight species. **(F)** Samples from (e) were digested with AlwNI, then separated on an agarose gel, and visualized by autoradiography. Red arrows indicate fragments enriched in phenol:chloroform purified samples. **(G)** Following lagging collapse, purified DNA samples were analyzed to detect end-joining products as for leading collapse in Fig. 1E-F. Colored arrows indicate end-to-end fusions of the same sizes indicated in Fig. 1E. **(H)** Phenol purified DNA samples from T=120 in (e) were treated with RuvC then separated on an agarose gel and visualized by autoradiography. RuvC treatment resolved most of the high molecular weight species, which indicated that they contained Holliday junctions **(I)** Quantification of DNA synthesis from Fig. 3B normalized to the maximum signal across all time points and conditions. Mean ± S.D., n = 3 independent experiments.

<u>**Figure S4: resolution of leading, but not lagging, collapse by extensive degradation**</u>

**(A)** Quantification of DNA synthesis from Fig. 4F normalized to the maximum signal across all time points and conditions. Mean ± S.D., n = 3 independent experiments. **(B)** Samples from T=120 in Fig. 4F were purified and restriction digested with AlwNI. Digested samples were then separated on an agarose gel and visualized by autoradiography. The collapsed arm fragment was readily detectable following MRE11 depletion, which indicated that degradation of the collapsed arm was blocked. **(C)** Quantification of collapsed arm products from (b). Mean ± S.D., n = 3 independent experiments. Data were analyzed by unpaired t-test. **(D)** Quantification of high molecular weight products (HMw) from Fig. 4F. Mean ± S.D., n = 3 independent experiments. **(E)** Quantification of full length products from (b). Mean ± S.D., n = 3 independent experiments. Data were analyzed by unpaired t-test. **(F)** Quantification of DNA synthesis from Fig. 4J normalized to the maximum signal across all time points and conditions. Mean ± S.D., n = 3 independent experiments. **(G)** Purified samples from T=120 in Fig. 4J were digested with XmnI and SacI, then separated on an agarose gel and visualized by autoradiography. **(H)** Quantification of DNA synthesis in the collapse region from (g) normalized to control fragment ‘Lin^1^’. Mean ± S.D., n = 3 independent experiments. Data were analyzed by one-way analysis of variance (ANOVA) and Tukey’s multiple-comparison method.

<u>**Figure S5: Analyses of bidirectional collapse in *Xenopus* egg extracts**</u>

**(A)** Quantification of DNA synthesis from Fig. 5B normalized to the maximum signal across all time points and conditions. Mean ± S.D., n = 4 independent experiments. **(B)** Quantification of linear products ‘Lin’ from Fig. 5B. Mean ± S.D., n = 4 independent experiments. **(C)** Quantification of collapse efficiency from Fig. 5B. Mean ± S.D., n = 4 independent experiments. Collapse efficiency was calculated as described in methods. **(D)** Bidirectional fork collapse was induced by replication of pSSB as in Fig. 5A. **(E)** DNA samples from T=120 minutes in (d) were purified by either phenol:chloroform extraction, to recover total DNA, or by spin column purification to preferentially exclude high molecular weight DNA. To investigate whether erroneous end-to-end fusions were formed, purified DNA was restriction digested with XmnI, AlwNI, or SapI, which should cut pSSB only once. Products of digestion were then separated on an agarose gel. Erroneous end-to-end fusions should have resulted in products greater than and smaller than the length of full length products (4661 bp), similar to Fig. 2E-F. However, exclusively linear products were formed in all conditions and few other species were generated, which indicated that replication of pSSB was largely accurate. **(F)** pSSB was replicated as in Fig. 4B lanes 6-10. DNA intermediates were purified, digested with AlwNI, then separated on an agarose gel. Replication and collapse intermediates were readily converted to full length linear products, which indicated that the ends were efficiently repaired. **(G)** pSSB was replicated as in Fig. 5B, lane 10, then treated with RuvC or left untreated. RuvC treatment did not have any effect on the abundance of high molecular weight species following bidirectional collapse (lanes 1-2), in contrast to leading collapse in Fig. 2D and lagging collapse in Fig. S3H. **(H)** DNA samples were prepared and purified as in (G) then treated with either AlwNI or topo II. High molecular weight products were resolved by AlwNI but not topo II, which indicated that they were not composed of catenanes. **(I)** Quantification of DNA synthesis from Fig. 5I normalized to the maximum signal across all time points and conditions. Mean ± S.D., n = 3 independent experiments. **(J)** Purified samples from T=120 of Fig. 5I were digested with XmnI and SacI to analyze DNA synthesis in the collapse region (as in Fig. 5E-G). Digested samples were separated on an agarose gel then visualized by autoradiography. **(K)** Quantification of DNA synthesis in the collapsed region from (j) normalized to control fragment ‘Lin^1^’. Mean ± S.D., n = 3 independent experiments. Data were analyzed by one-way analysis of variance (ANOVA) and Dunnett’s multiple-comparison method. No detectable effect on collapse region DNA synthesis was observed, which indicated that completion of DNA synthesis occurred independently of RAD51.

<u>**Figure S6: Analyses of nCas9 induced collapse in *Xenopus* egg extracts**</u>

**(A)** Quantification of DNA synthesis from Fig. 6B. Mean ± S.D., n = 3 independent experiments. **(B)** Quantification of nicked plus supercoiled circular monomers (CMs) from Fig. 6B. Mean ± S.D., n = 3 independent experiments. **(C)** Quantification of σ structures plus linear products (collapsed molecules) from Fig. 6B. Mean ± S.D., n = 3 independent experiments. **(D)** Quantification of θ structures from Fig. 6B. Mean ± S.D., n = 3 independent experiments. **(E)** Quantification of collapse efficiency from Fig. 6B. Mean ± S.D., n = 3 independent experiments. Collapse efficiency was calculated as described in methods. **(F)** Leading collapse was induced using nCas9 as in Fig. 6B lanes 6-10 in the presence of either vehicle, BRC peptide (BRC WT), or BRC mutant (BRC Mut) peptide. DNA samples were separated on an agarose gel and visualized by autoradiography**. (G)** Quantification of high molecular weight products (HMw) from (f). Mean ± S.D., n = 3 independent experiments. **(H)** Quantification of DNA synthesis from (f) normalized to the maximum signal across all time points and conditions. Mean ± S.D., n = 3 independent experiments. **(I)** Purified DNA samples from T=120 in (f) were analyzed as in Fig. 6F to monitor synthesis in the collapse region. **(J)** Synthesis in the collapse region was quantified from (I) as in Fig. 6G. Mean ± S.D., n = 3 independent experiments. Data were analyzed by one-way analysis of variance (ANOVA) and Dunnett’s multiple-comparison method. **(K)** Bidirectional collapse was induced using nCas9 as in Fig. 6B lanes 16-20 in the presence of either vehicle, BRC peptide (BRC WT), or BRC mutant (BRC Mut) peptide. DNA samples were separated on an agarose gel and visualized by autoradiography**. (L)** Quantification of high molecular weight products (HMw) from (k). Mean ± S.D., n = 3 independent experiments. **(M)** Quantification of DNA synthesis from (k) normalized to the maximum signal across all time points and conditions. Mean ± S.D., n = 3 independent experiments. **(N)** Purified DNA samples from T=120 in (k) were analyzed as in Fig. 6F to monitor synthesis in the collapse region. **(O)** Synthesis in the collapse region was quantified from (N) as in Fig. 6G. Mean ± S.D., n = 3 independent experiments. Data were analyzed by one-way analysis of variance (ANOVA) and Dunnett’s multiple-comparison method. **(P)** Leading collapse and bidirectional collapse were induced by replication of pCRISPR as in Fig. 6A lanes 5, 10, 15 and 20. Purified DNA products were treated with RuvC then separated on an agarose gel and visualized by autoradiography. High molecular weight products (HMw) are mostly sensitive to RuvC for leading collapse (lanes 1-2) but mostly insensitive for bidirectional collapse (lanes 3-4) as observed for collapse at stabilized SSBs (Fig. 2D, Fig. S5G). **(Q)** Purified DNA was obtained as in (p) then treated either AlwNI or TopoII, separated on an agarose gel and visualized by autoradiography. High molecular weight (HMw) species were resolved by digestion with AlwNI but not topo II, which indicated that they were not composed of catenanes, as observed for collapse at stabilized SSBs (Fig. S2H, Fig. S5H). **(R)** Quantification of nicked circular monomers (nCM) from (Fig. 6B). Mean ± S.D., n = 3 independent experiments. **(S)** Lagging collapse was induced by replication of pCRISPR as depicted in (Fig. 6a) but with guides targeting the lagging strand template. Replication samples were separated on an agarose gel and visualized by autoradiography. **(T)** Leading collapse was induced using nCas9 as in Fig. 6A but using pCRISPR^1X^ that contained only a single target sequence. Replication samples were separated on an agarose gel and visualized by autoradiography. **(U)** Quantification of θ structures from (t). Mean ± S.D., n = 6 independent experiments. **(V)** Quantification of nicked plus supercoiled circular monomers from (t). Mean ± S.D., n = 6 independent experiments. **(W)** Quantification of collapse efficiency from (t). Mean ± S.D., n = 6 independent experiments. Collapse efficiency was calculated as described in methods.

<u>**Figure S7: Analyses of SSBs in *Xenopus* egg extracts**</u>

**(A)** Plasmid DNA containing a deoxyuridine (pdU^LAG^) or deoxythymidine control site (pdT^CTRL^) was treated with uracil DNA glycosylase ‘UDG’ to induce abasic site (AP site) formation. The resulting products were then treated with APE1 to use APE1-induced nicking as a readout for the presence of an AP site. DNA products were separated on an agarose gel and visualized by SYBR Gold staining. APE1 treatment nicked essentially all supercoiled circular monomers (scCMs) for pdU^LAG^ but had no such effect on pdT^CTRL^. Thus, AP sites were efficiently generated from pdU^LAG^, as previously described^115^. **(B)** To assess the stability of AP sites in Xenopus egg extract, pdT^CTRL^ and pdU^LAG^ were treated with or without uracil DNA glycosylase ‘UDG’ then incubated with or without LacR (to stabilize the AP site, as in Fig. 7A) prior to incubation in High Speed Supernatant (HSS) of *Xenopus* egg extract that supports licensing but not initiation of DNA replication. Products were separated on an agarose gel then visualized by SYBR Gold staining. Following UDG treatment, pdU^LAG^ became extensively nicked (lanes 7-9), and this was not detected in the absence of LacR (lanes 10-18) or for pdT^CTRL^ (lanes 1-3). Thus, LacR-bound AP sites were converted by *Xenopus* egg extract to stable SSBs. Notably, pdU^LAG^ underwent a low amount of nicking even without UDG treatment (lanes 4-6), which indicated that deoxyuridine itself was sufficient to give rise to a stable SSB, likely by conversion to an AP site by UDG present in *Xenopus* egg extract. **(C)** Quantification of total DNA synthesis from Fig. 7B normalized to the maximum signal across all time points and conditions. Mean ± S.D., n = 3 independent experiments. **(D)** Structure of the PARP1/2 inhibitor Talazoparib (PARPi) that efficiently inhibits PARP activity in *Xenopus* egg extracts^77^. **(E)** LacR-bound plasmid pdT^CTRL^ was replicated as in Fig. 7D in either the presence or absence of PARPi. Replication products were then separated on an agarose gel and visualized by autoradiography. **(F)** Quantification of σ structures (σs) from (e). Mean ± S.D., n = 3 independent experiments. PARPi did not appreciably increase the abundance of σs, which indicated there was no measurable effect on fork collapse. **(G)** Quantification of high molecular weight products (HMw) from (e). Mean ± S.D., n = 3 independent experiments. PARPi did not increase formation of HMw products, consistent with lack of collapse induced by PARPi. **(H)** pSSB^1X^ was replicated as in (Fig. 7i), except TetR was omitted from the reaction. Replication products were then separated on an agarose gel and visualized by autoradiography. **(I)** Quantification of σ structures (σs) from (h). Mean ± S.D., n = 3 independent experiments. PARPi did not appreciably increase the abundance of σs, which indicated there was no measurable effect on fork collapse. **(J)** Quantification of high molecular weight products (HMw) from (h). Mean ± S.D., n = 3 independent experiments. PARPi did not increase formation of HMw products, consistent with lack of collapse induced by PARPi. **(K)** Fork collapse was induced by replication of plasmid pSSB^1X^ in the presence of LacR, TetR, and PARPi and the presence of either vehicle, BRC peptide (BRC WT), or BRC mutant (BRC Mut) peptide. **(L)** Samples from (k) were separated on an agarose gel and visualized by autoradiography. **(M)** Quantification of high molecular weight products from (l). Mean ± S.D., n = 3 independent experiments. Collapse was readily detectable but a high degree of experiment-to-experiment variability in the level of collapse precluded any definitive conclusion from the data. **(N)** To account for the experimental variability in (m), quantification for each individual experimental replicate was normalized to the 120 minute time point in the vehicle condition. The resulting quantification of normalized high molecular weight products (HMw) from (l) showed that BRC^WT^ severely inhibited HMw formation and this was diminished by BRC^Mut^, indicating that generation of HMw species was at least partially dependent on RAD51. Mean ± S.D., n = 3 independent experiments.

## Methods

### Statistics and probability

For replication in *Xenopus* egg extract, the number of experimental repeats and the statistical analysis applied are provided in the corresponding figure legend. Statistical analyses were performed using GraphPad Prism v10. Statistical significance is indicated in figures using *s, where *, **, ***, and **** represent p-values of ≤ 0.05, 0.005, 0.0005, and 0.0001 respectively.

### *Xenopus* egg extract

*Xenopus* egg extracts were prepared from wild-type *Xenopus laevis* male and female frogs (Xenopus1) as previously described^126^. Animal protocols were approved by the Vanderbilt Division of Animal Care and the Institutional Animal Care and Use Committee.

### Plasmid construction

Commercially available plasmid pET-28b(+) (Novagen), referred to as ‘ctrl’ and given the identifier pJD1, was used as an internal loading control in indicated experiments. Plasmids pSSB^LEAD^, pSSB^LAG^, pSSB^1X^, pCRISPR, pdT^CTRL^, pdU^LAG^, and pAP are modified derivatives of pJD161, which has been described previously^98^. In brief, pJD161 contains 50x tandem repeats of the *lac* operator (“*lac*O”) sequence that collectively form a ∼1600 bp *lac*O array. To create pSSB^LEAD^ and pSSB^LAG^ blunt-ended DNA duplex JDD27 (5’-TCTCTATCACTGATAGGGAATGCTCTC TATCACTGATAGGGAATGCTCTCTATCACTGATAGGGAATGCTCTCTATCACTGATAGGGAAT GCTCTCTATCACTGATAGGGAATGCTCTCTATCACTGATAGGGA-3’)^Top^ ^Strand^/(5’-TCCCTATCA GTGATAGAGAGCATTCCCTATCAGTGATAGAGAGCATTCCCTATCAGTGATAGAGAGCATTCCCTATCAGTGATAGAGAGCATTCCCTATCAGTGATAGAGAGCATTCCCTATCAGTGATAGAGA-3’)^Bottom^ ^Strand^ was cloned into pJD161 that had been linearized with PsiI (NEB). The same steps were performed for pSSB^1X^, except DNA duplex JDD26 (5’-TCCCTATCAGTGATAGAGATC CCTATCAGTGATAGAGA-3’)^Top^ ^Strand^/(5’-TCTCTATCACTGATAGGGAATGCTCTCTATCACTGAT AGGGA-3’)^Bottom^ ^Strand^ was utilized instead. JDD27 contained 5x tandem Nb.BsmI restriction sites flanked by a single *tetracycline* operator (“*tetO*”) sequence on either side while JDD26 contained only 1x Nb.BsmI restriction site that was *tet*O-flanked. Blunt-end ligation of linearized pJD161 with JDD26 or JDD27 allowed for insertion of the DNA duplexes in both forward and reverse orientations. The resulting ligation products were transformed into DH5α cells. Constructs were chosen that contained JDD26 and JDD27 in both orientations such that digestion with Nb.BsmI would nick either the top (pSSB^LAG^) or bottom (pSSB^LEAD^ and pSSB^1X^) strand.

To create pCRISPR, DNA oligonucleotides JDO143 (5’-CCAAACTGGAACAACACTCA ACCCTATCTCGGGTGACATACGAGTCTTACCAAACTGGAACAACACTCAACCCTATCTCGGC CAGACAGTGGACTCTGCCAAACTGGAACAACACTCAACCCTATCTCGGAGGCAGAATCGCC CGTACCAAACTGGAACAACACTCAACCCTATCTCGGTTACATTCTCACCGTCGTTA-3’) and JDO144 (5’-TAACGACGGTGAGAATGTAACCGAGATAGGGTTGAGTGTTGTTCCAGTTTGGTACGGGCGATTCTGCCTCCGAGATAGGGTTGAGTGTTGTTCCAGTTTGGCAGAGTCCACTGTCTGGCCGAGATAGGGTTGAGTGTTGTTCCAGTTTGGTAAGACTCGTATGTCACCCGAGATAGGGTTGAGTGTTGTTCCAGTTTGG-3’) were annealed together and cloned into pJD161 that had been linearized using PsiI (NEB). Constructs were chosen that contained JDO144 in the top strand and JDO143 in the bottom strand in a 5’ to 3’ orientation. The resulting plasmid, pCRISPR, contained 5x tandem repeats of the target DNA sequences (5’-GGTTGAGTGTTG TTCCAGTT-3) and (5’-AGATAGGGTTGAGTGTTGTT-3’), which could be targeted by nCas9^D840A^ to generate SSBs in the leading and lagging strands respectively.

Plasmids pdU^LAG^ and pdT^CTRL^ were prepared similarly to those previously described^115^. In brief, DNA oligonucleotides oMC13 (5’-ATTATCTAGACCTCAGCTTGTGAGCGGATAACAAGCATTTGTGA GCGGATAACAACCTCAGCTAGCTTA-3’) and oMC14 (5’-TAAGCTAGCTGAGGTTGTTATC CGCTCACAAATGCTTGTTATCCGCTCACAAGCTGAGGTCTAGATAAT-3’) were annealed together to form a DNA duplex that contained two tandem Nt.BbvCI nicking sites which, upon digestion with Nt.BbvCI (NEB), would allow for insertion of custom oligonucleotide sequences. Duplexed oMC13/oMC14 was cloned into pJD161 that had been linearized using PsiI (NEB), and constructs were chosen that contained oMC13 in the top strand in a 5’ to 3’ orientation. The resulting plasmid, termed pMC3, was subsequently used to prepare modified plasmids pdU^LAG^, which contained a deoxyuridine (“dU”) site, and pdT^CTRL^, which lacked the dU site and served as a control plasmid for pdU^LAG^. Plasmid pAP was prepared from pdU^LAG^ and its preparation is described below.

### Protein purification

Tetracycline repressor protein (“TetR”) and biotinylated lac repressor protein (“LacR”) were expressed in T7 express Escherichia coli (NEB) and purified as previously described^107^.

### Preparation of damaged plasmid templates

To generate SSB-containing plasmid DNA, pSSB^LEAD^, pSSB^LAG^, and pSSB^1X^, 30 μg of plasmid DNA was digested with 75 U of nicking enzyme Nb.BsmI (NEB) in 1x NEBuffer r3.1 (NEB) for 1 hour, 45 min at 37°C. Nicked DNA was then resolved on a 0.9% agarose gel at 5 V/cm and stained with SYBR Gold (Invitrogen). The nicked DNA band was excised using a blue light transilluminator and subsequently purified using electroelution. Electroelution was performed by placing the excised gel slice in SnakeSkin dialysis tubing (3.5 kDa molecular weight cut-off, 35 mm diameter, Thermo Scientific) that contained 1 mL of 1X TBE supplemented with bovine serum albumin (final concentration, 0.3 mg/mL). The gel slice was then electrophoresed at 5 V/cm for 1 hour, 30 min. The slice was subsequently discarded while the purified DNA, which migrated into the buffer, was left in the dialysis tubing and was dialyzed overnight at 4°C in 1 L 10 mM Tris (pH 8.0). Dialyzed DNA was then collected and concentrated using Amicon centrifugal concentrators (100 kDa molecular weight cut-off, 0.5 mL volume). To estimate DNA concentration, purified SSB-containing plasmids were resolved alongside a DNA standard that had been purified by phenol:chloroform:isoamyl alcohol extraction (25:24:1) followed by ethanol precipitation (70% (vol/vol) final) in the presence of sodium acetate (270 mM final) and glycogen (1% final). SSB-containing plasmid DNAs were brought to a final concentration of 225 ng/μl and were stored at -20°C until used in experiments.

To generate plasmids pdT^CTRL^ and pdU^LAG^, 20 μg of pMC3 was nicked with 75 U of Nt.BbvCI in 1x rCutsmart Buffer (NEB) for 1 hour at 37°C, and nicked products were purified by spin column purification according to the manufacturer’s instructions (PCR Purification Kit, Qiagen). Oligonucleotide oMC16 (5’-TCAGCTTGTGAGCGGATAACAAGCA<u>/ideoxyU/</u>TTG TGAGCGGATAACAACC-3’), which contained a dU site, and oMC15 (5’-TCAGCTT GTGAGCGGATAACAAGCA<u>T</u>TTGTGAGCGGATAACAACC-3’), a control sequence that lacked the dU site, were annealed into nicked pMC3 to create plasmids pdU^LAG^ and pdT^CTRL^ respectively. To accomplish this, 8.5 μg of pMC3 was combined with 100-fold molar excess of oMC15 or oMC16, heated to 70°C for 5 min, and then cooled to 25°C at a rate of -1°C per minute in a thermocycler. The resulting annealed products were then ligated with 1200 U of T4 DNA ligase in 1x T4 Ligase buffer supplemented with 5 mM ATP overnight at room temperature. To remove any excess unligated plasmid and/or oligos, ligation products were resolved on a 0.8% TAE-agarose gel (+0.3 ug/mL EtBr). The supercoiled fraction (i.e. ligated plasmids) was excised from the gel and purified using a gel extraction kit (Qiagen). The final products were buffer exchanged into 10 mM Tris-HCl, pH 8 using Amicon centrifugal concentrators (100 kDa molecular weight cut-off, 0.5 mL volume), and then brought to a concentration of 300 ng/μl. To generate the abasic site in pdU^LAG^, and thus form plasmid pAP, 675 ng plasmid DNA was treated with 2.25 U of uracil DNA glycosylase (“UDG”; NEB) in 1x UDG Reaction Buffer (NEB) for 1 hour at 37°C in a 3 μL reaction volume. These steps were performed immediately prior to replication in extracts (see section below). Additionally, pdT^CTRL^, which did not contain a dU site, also underwent UDG treatment and served as a control in experiments.

To generate 1x or 5x tandem SSBs in DNA using nCas9, pCRISPR^1X^ (pJD161) or pCRISPR was incubated in licensing in extract (see section below) along with assembled CRISPR-Cas9 RNP (“ribonucleoprotein”, see section below).

### Assembly of CRISPR-Cas9 Ribonucleoprotein Complex (“RNP”)

The CRISPR-Cas9 RNP complex was assembled by incubation of guide RNA (“gRNA”) with either Alt-R S.p. dCas9 Protein V3 (IDT) or Alt-R S.p. Cas9 H840A Nickase V3 (IDT). gRNA was prepared by mixing Alt-R CRISPR-Cas9 tracrRNA, ATTO550 (IDT) with 10-fold molar excess Alt-R CRISPR-Cas9 crRNA in nuclease-free duplex buffer (IDT). Two crRNAs, scRNA1 (5’-GGUUGAGUGUUGUUCCAGUU -3’) and scRNA2 (5’-AACAACACUCAACCCUAUCU-3’) were used to assemble guide RNA that targeted leading or lagging strands respectively. The crRNA-tracrRNA mixtures were heated to 95°C for 5 minutes then cooled to room temperature on the benchtop for 1 hour. Prepared guide RNA was stored at -20°C until used in experiments. Immediately prior to experiments, the CRISPR-Cas9 RNP complex was formed by incubating 25 μM of Cas9 enzyme with 2 μM of gRNA in 1X Cas9 dilution buffer (15 mM KCl, 3 mM HEPES, pH 7.5), in a 2 μL reaction volume. This mixture was incubated at room temperature for at least 20 minutes prior to use in experiments.

### DNA replication in Xenopus egg extracts

Control plasmid (pJD1) was fully replicated in extracts prior to induction of fork collapse. To accomplish this, activated High Speed Supernatant (“HSS”) was prepared by incubating HSS with ATP regenerating system (“ARS”; 20 mM phosphocreatine, 2 mM ATP, and 5 ng/μl creatine phosphokinase) and nocodazole (3 ng/μl) for 5 min at room temperature. Plasmid DNA was then licensed by addition of 0.1 volumes of pJD1 (100 ng/μl) to 0.9 volumes activated HSS. The “licensing mix” was incubated for 20 min at room temperature. Activated Nucleoplasmic Extract (“NPE”) was prepared by incubating NPE with ARS, dithiothreitol (“DTT”; final concentration, 2 mM) and [α-^32^P]dATP (final concentration, 350 nM). The “NPE mix” was then diluted with 1x egg lysis buffer (“ELB”; 250 mM sucrose, 2.5 mM MgCl_2_ 50 mM KCl, 10 mM HEPES, pH 7.7) to achieve a final NPE concentration of 45% (vol/vol). Replication was initiated by adding 0.1 volume of licensing mix to 0.9 volumes of 45% NPE mix. Replication of the control plasmid was allowed to proceed for 40 min before 0.05 volumes of the reaction was added to 0.95 volumes of fresh NPE mix that had been diluted to 65% in 1X ELB. This “control-NPE mix”, which contained fully replicated pJD1, was then used in fork collapse experiments. Re-replication of pJD1 did not occur because NPE contains a high concentration of factors which inhibit origin licensing^99^.

Fork collapse was induced by replicating SSB- or AP site-containing plasmids in *Xenopus* egg extract. For experiments that used either pSSB^LEAD^, pSSB^LAG^, or pSSB^1X^ as template, 0.29 volumes of pSSB (225 ng/μl) (see section above) was incubated with 0.36 volumes of either TetR (100 μM) or TetR buffer and 0.36 volumes of either LacR (32 μM) or LacR buffer for 1 hour at room temperature. These steps were performed immediately prior to licensing in activated HSS. To license pSSB plasmids, 0.2 volumes of repressor-bound plasmid was added to 0.8 volumes of activated HSS for a final DNA concentration of 13 ng/μl. These “licensing mixes” were then incubated for 20 min at room temperature. Replication of pSSB was initiated by addition of 1 volume of licensing mix to 2 volumes of control-NPE mix. For reactions that utilized TetR to stabilize SSBs, 0.05 volumes of TetR protein (100 μM) was added to 0.95 volumes of the control NPE mix prior to replication. Where indicated, BRC peptide was added to reactions at a final concentration of 18 μM. The PARP inhibitor talazoparib (Selleckchem) was dissolved in DMSO and added to reactions at a final concentration of 80 μM.

Replication of plasmids pdT^CTRL^ and pAP was performed essentially as outlined above for pSSB plasmids except that 0.64 volumes of plasmid DNA (98 ng/μl, see section above) was incubated with 0.36 volumes of LacR (32 μM) for 1 hour at room temperature prior to licensing in activated HSS. All subsequent steps (i.e., licensing, initiation, etc.) were the same.

For experiments that used pCRISPR or pJD161 to induce collapse, 0.57 volumes of plasmid DNA (115 ng/μl) were bound with 0.43 volumes of either LacR (32 μM) or LacR buffer for 1 hour at room temperature. CRISPR-Cas9 RNP complexes were assembled (see section above), diluted 1.75-fold in H_2_O, and 0.57 volumes of diluted RNP complex was incubated with 0.43 volumes of LacR (32 μM) or LacR buffer for 1 hour at room temperature. Plasmids pCRISPR or pJD161 were then licensed by incubation of 0.1 volumes of repressor-bound plasmid with 0.8 volumes of activated HSS for 20 minutes at room temperature. After 20 minutes, 0.1 volumes of the prepared CRISPR-Cas9 RNP was added to the licensing mix, and licensing was allowed to proceed for an additional 10 minutes for a total incubation time of approximately 30 minutes. As for plasmids pSSB, dT^CTRL^, and pAP, replication of pCRISPR was initiated by addition of 1 volume of licensing mix to 2 volumes of “control” NPE mix.

At the indicated time points, reactions were sampled into either 20 volumes of replication stop solution (8 mM EDTA, 0.13% phosphoric acid, 10% ficoll, 5% SDS, 0.2% bromophenol blue, 80 mM Tris pH 8.0), which stops replication reactions and serves as a DNA loading dye, or 20 volumes of extraction stop solution (1% SDS, 25 mM EDTA, 50 mM Tris-HCl pH 7.5), which also stops reactions but is compatible with downstream processing (i.e., DNA purification). Stopped reactions were treated with RNase A (final concentration, 174 ng/μl) followed by Proteinase K (final concentration, 1.6 mg/mL). Samples collected in replication stop were subsequently analyzed by agarose gel electrophoresis at 5 V/cm. Samples collected in extraction stop were subsequently purified by either column purification (Monarch PCR & DNA Cleanup Kit, NEB) or phenol:chloroform extraction followed by ethanol precipitation (70% (vol/vol) final) in the presence of sodium acetate (270 mM final) and glycogen (1% final). Purified DNA samples were resuspended in 6 μl of 10 mM Tris-HCl (pH 8.0).

Experiments conducted in Fig. 1-5 and Fig. S1-5 used either pSSB^LEAD^ or pSSB^LAG^ as the template for replication. Experiments conducted in Fig. 6B-D,F-G and Fig. S6A-S used pCRISPR as template while experiments in Fig. 6h-i and Fig. 6ST-W used pJD161, alternatively referred to as pCRISPR^1X^, as template. For experiments shown in Fig.7b-c,e-h and Fig. 7SC, pAP was used as template while pdT^CTRL^ was used as template in experiments shown in Fig. S7E-G. Lastly, pSSB^1X^ was used as template in experiments conducted in Fig. 7j-m and Fig. S7L-N. Control plasmid (pJD1) was included as a loading control in almost all experiments with the following exceptions, for which it was omitted: Fig. S3G, Fig. 4E-H, and Fig. S4A-E.

### Antibodies and Peptides

Antibody targeting *Xenopus* MRE11 was raised by New England Peptide by immunizing rabbits with Ac-CDPFKKSGPSRRGRR-OH. BRC peptides were prepared as previously described^127^.

### Immunodepletion

Immunodepletion of *Xenopus* egg extract was performed as previously described^128^, with slight modifications. In brief, Protein A coupled magnetic beads (Dynabeads Protein A 30 μg/μL) were bound with 0.5 μg of either control IgGs or anti-Mre11 antibody per 1 μg of magnetic beads. For each round of depletion, 1.29 volumes of antibody-bound beads were incubated with 0.5 volumes of HSS or 1 volume of NPE for 20 min at room temp with end-over-end rotation. This was repeated once for HSS, for a total of 2 rounds of depletion, and twice for NPE, for a total of 3 rounds of depletion. Depleted extracts were harvested and subsequently used for DNA replication (as described above).

### Analysis of replication and repair intermediates

To monitor replication, samples that had been collected into replication stop were separated on a 1% agarose gel at 5 V/cm. Radiolabeled DNA was detected by phosphorimaging, which measured the incorporation of radiolabeled nucleotides, and DNA signal was measured using ImageJ. To measure DNA synthesis, individual whole lane signals were first normalized to the loading control present in each lane. The normalized whole lane signals were then expressed relative to the maximum whole lane signal across all time points and all conditions (AU). Due to nicking of the internal loading control by nCas9, DNA synthesis was not normalized to loading control for CRISPR-Cas9 experiments (Fig. 6 and Fig. S6). DNA synthesis was also not normalized to loading control for MRE11 depletion experiments (Fig. 4 and Fig. S4) due to omission of the loading control in order to more clearly visualize circular monomers in these experiments.

The abundance of individual DNA species was expressed as a percentage of whole lane signal (%). Collapse efficiency in single fork conditions was calculated as [1 – (θ% in collapse conditions/θ% in control conditions)]x100 for each timepoint. The principle behind this formula is that replication fork structures (θs) are reduced in a manner that is directly proportional to the frequency of collapse due to conversion of θs to σs. For converging fork experiments in which plasmids pSSB^LEAD^, pSSB^LAG^, or pCRISPR were used template, collapse efficiency was calculated as [(([(nCM% + scCM%) / Lin%] – 1)/2) + 1]^-1^ x 100%. The principle behind this formula is that for converging forks each bidirectional collapse event should generate a nicked or supercoiled circular monomer as well as a linear molecule while completion of DNA synthesis without collapse will generate two nicked or supercoiled Circular Monomers. For converging fork experiments in which plasmid pJD161 (i.e., pCRISPR^1X^) was used template, the aforementioned formula was not used to calculate collapse efficiency, as the linear band generated from collapse migrated at the same position as supercoiled control plasmid. Collapse efficiency was instead calculated using a less direct approach, as follows. First, circular monomers were normalized to the control: CMnorm = (nCM%+scCM% in collapse conditions)/(maximum nCMs%+scCM% in control conditions). Second, CMnorm was used to calculate collapse efficiency across all time points by applying the following formula: [1 – ([CMnorm – 0.5]/0.5)] x100%. The principle behind this formula is that each collapse event should lead to loss of one Circular Monomer out of the two that would be formed if replication proceeded without collapse.

### Analysis of DNA synthesis in the collapse region

To monitor DNA synthesis in the collapse region, purified DNA samples were digested with 0.4 U/μl XmnI and 0.4 U/μl SacI in 1x rCutSmartBuffer (NEB) for 1 hour at 37°C. Digested products were then separated under either native or denaturing agarose gel conditions. For separation under native conditions, 6x native loading buffer (Ficoll, EDTA, SDS, bromophenol blue, and Tris-HCl (1M, pH 8)) was added to samples at a final concentration of 1x before separation on a 1% agarose gel at 5 V/cm. Radiolabeled DNA was subsequently detected by phosphorimaging. For separation under denaturing conditions, digests were stopped by addition of EDTA to a final concentration of 30 mM before addition of 6x alkaline loading buffer (Ficoll, EDTA, xylene cyanol, bromocresol green, and NaOH (10N)) to a final concentration of 1x. Digests were then separated on a 1.5% alkaline gel at 1.5 V/cm. The gel was then neutralized with gentle agitation in 7% trichloroacetic acid, and radiolabeled DNA was detected by phosphorimaging.

DNA synthesis in the collapse region was calculated using the following formula, where: a = signal of the collapsed fragment; and b = signal of pJD1 control fragment 1; as [((a / b) in the collapse condition) / (average (a / b) in no collapse control i.e. Tet buffer or vehicle conditions)] x 100%. For single fork collapse experiments that used either pSSB^LEAD^ or pSSB^LAG^ as template, “expected no repair” was calculated as ([(747/2212) x fraction of molecules that collapsed] + fraction of molecules that did not collapse) x DNA synthesis% within the collapse region in control conditions. In the above formula, the value (747/2212) corresponds to the amount of nascent DNA synthesis within the collapsed region, assuming no repair. To adjust “expected no repair” for differences in replication efficiency, the calculated value was then multiplied by (DNA Synthesis (AU) at T=120 in collapsed conditions / DNA Synthesis (AU) at T=120 in control conditions). For nCas9 induced single fork collapse experiments that used template plasmids pCRISPR or pJD161, alternatively referred to as pCRISPR^1X^, the same calculations as above were used except (724/2336) and (625/1944) were used respectively as the expected nascent synthesis within the collapsed region assuming no repair of collapsed ends. For collapse under converging fork conditions, the same calculations were applied, except 0.5 was used as the expected signal assuming no repair because the parental strand lacking a SSB was fully replicated.

### Enzymatic analysis of replication and repair intermediates

Where indicated, purified DNA was subjected to the following enzymatic digestions: single digestion with 0.8 U/μl of AlwNI, XmnI, or SapI in 1X rCutSmartBuffer (NEB) for 1 hour at 37°C, double digestion with 0.4 U/μl SacI and 0.4 U/μl KpnI in 1x rCutSmartBuffer (NEB) for 1 hour at 37°C, single digestion with 0.12 U/μl of hTOP2α in 1X Topoisomerase II Assay Buffer (Topogen) for 15 min at 37°C, or single digestion with 0.5 nM to 1.5 nM RuvC in 1X NEBuffer 2.1(NEB) supplemented with DTT (final concentration, 1mM) and Tris, pH 8.0 (final concentration, 40mM) for 1 hour at 37°C. hTOP2α digestion was stopped by addition of 0.5 volumes TopSTOP solution (3% SDS, Proteinase K 2 mg/mL). RuvC digestion was stopped by addition of 0.2 volumes RuvCSTOP solution (3% SDS, 240 mM EDTA, Proteinase K 3 mg/mL). The radiolabeled digestion products were then separated on a 1% agarose gel at 5 V/cm and detected by phosphorimaging.

### End-labeling of DNA

Where indicated, digested DNA products were run alongside a radiolabeled DNA ladder. Radiolabeled ladder was generated by treating 2 μg DNA ladder (NEB) with 1 U T4 Polynucleotide Kinase (“PNK”, NEB) in 1X PNK Reaction Buffer (NEB) supplemented with 1.33 mM [γ-32P]ATP in a 20 μL total volume. The reaction was allowed to proceed for 45 mins at 37°C, and then purified using the Monarch PCR & DNA Cleanup Kit (NEB) according to the manufacturer’s instructions.

### Stability of AP sites in extract

To test the stability of abasic sites in extract, pdT^CTRL^ (225 ng/μl) and pdU^LAG^ (225 ng/μl) were treated with or without 0.75 U/μl uracil DNA glycosylase (“UDG”, NEB) in 1x UDG Reaction Buffer (NEB) for 1 hour at 37°C. Plasmid DNAs were then diluted 2.3-fold in H_2_O and 0.64 volumes of diluted plasmid was incubated with 0.36 volumes LacR (32 μM) or LacR buffer for 1 hour at room temperature. 0.2 volumes of plasmid DNA were then incubated in 0.8 volumes of activated HSS (described above). At the indicated time points, 2 volumes of reaction were sampled into 98 volumes of stop solution (1% SDS, 20 mM EDTA). Stopped reactions were treated with RNAse A (40 μg/mL) for 30 minutes at 37°C followed by treatment with Proteinase K (194 μg/mL) for 1 hour at 37°C. Samples were then resolved on a 1% agarose gel containing 0.3 μg/mL ethidium bromide.

## Conflict of interest

The authors declare no competing interests.

## Data availability

All data supporting the findings of this study are available within the manuscript and its supplement. Raw data are available upon request.

## Acknowledgements

JMD was supported by NIH grant R35GM128696. DC was supported by R01ES030575. DTL was supported by R35GM119512. SCC was supported by T32CA009582 and T32GM139800. MTC was supported by NIH grant T32ES007028 and F32GM148024. The project received supplemental support from P50CA098131. We thank Savannah Weeks Pollenz for helpful feedback on the manuscript.

## Author contributions

JMD conceived of and supervised the project. SCC performed all work except where noted otherwise. JMD, DC, and MTC made the initial observation that AP sites could induce efficient fork collapse. MTC generated pdU and pdT for AP site analysis. SND generated the pCRISPR-5X plasmid. RJM made the initial observation that PARP inhibition stabilizes a TetR-bound SSB. DTL purified and validated BRC peptides. The manuscript was written by JMD and SCC.

## Notes

### Competing Interest Statement

The authors have declared no competing interest.

