## Supplementary figures and images for "Resolution of collapsed forks is separate from completion of DNA synthesis"

### Supplemental Figures

#### Supp Figure 1

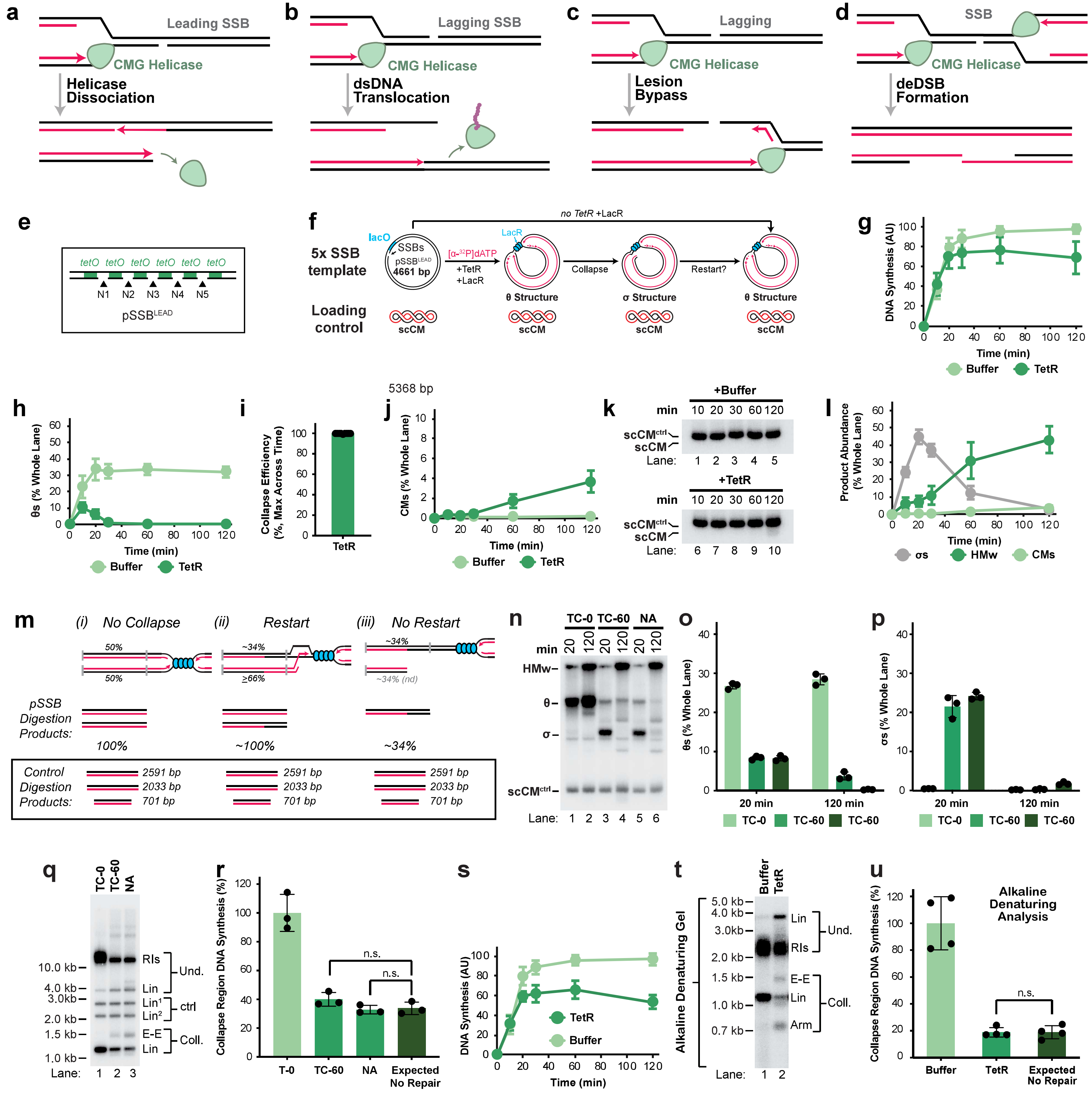

#### Supp Figure 2

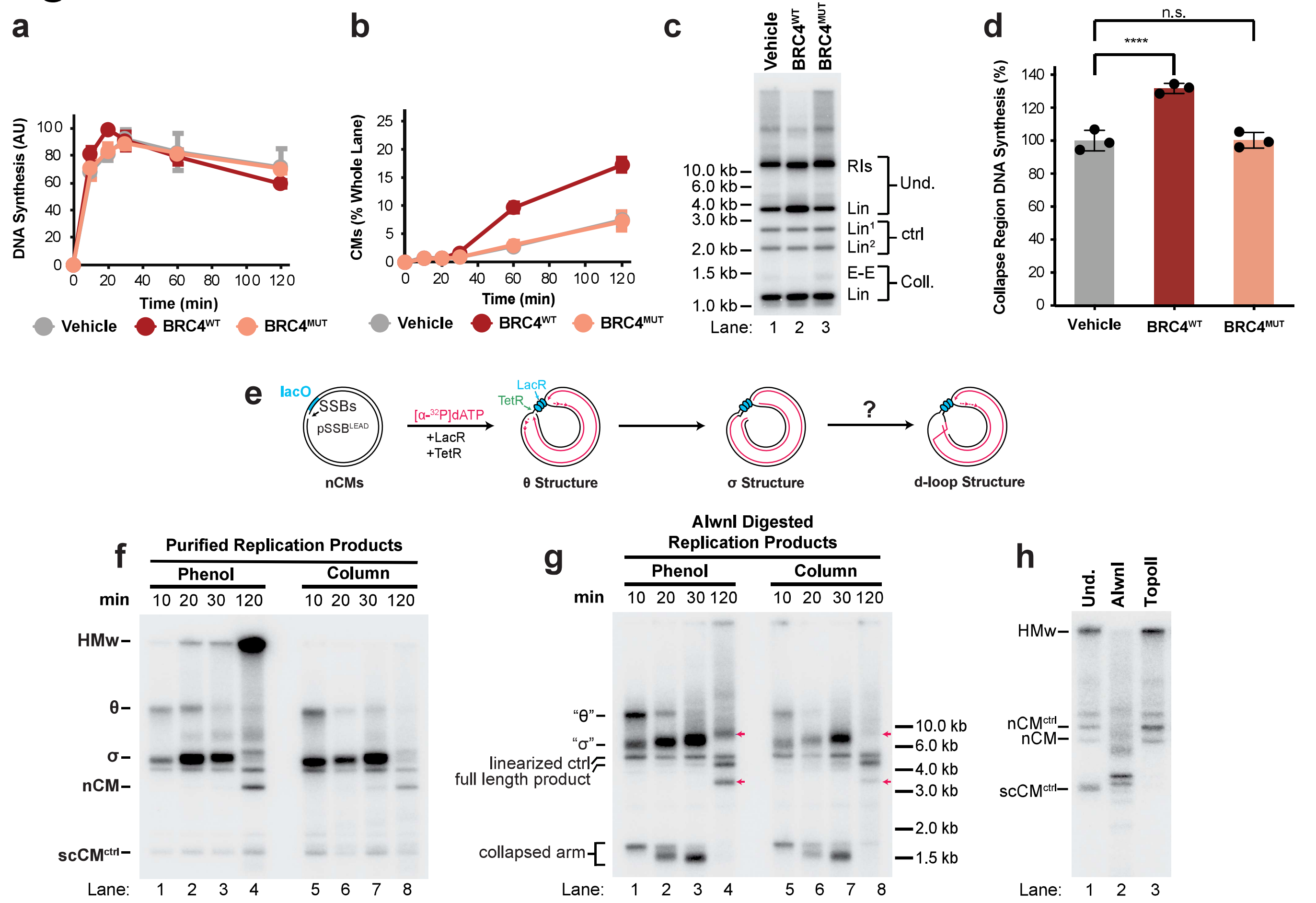

#### Supp Figure 3

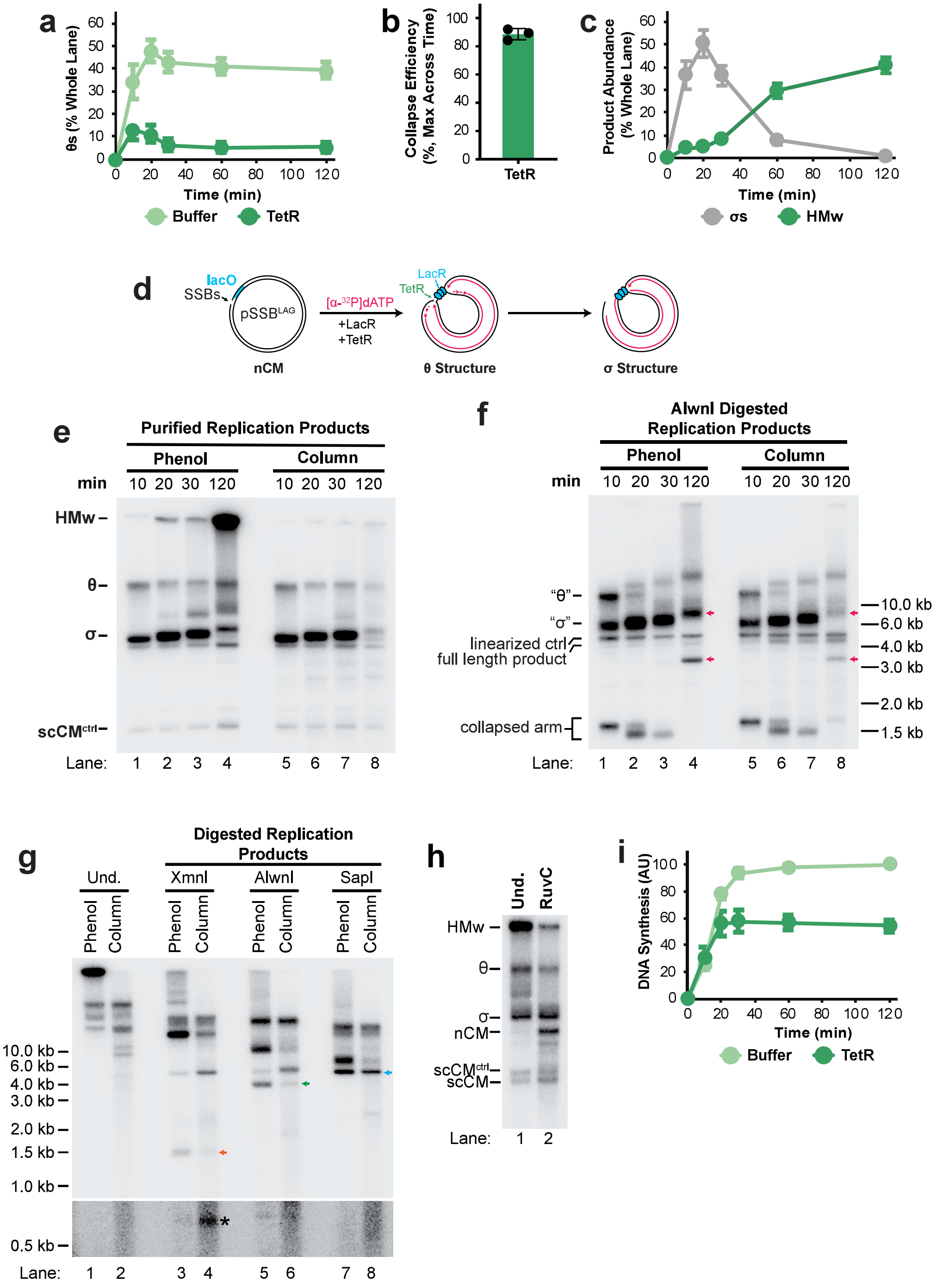

#### Supp Figure 4

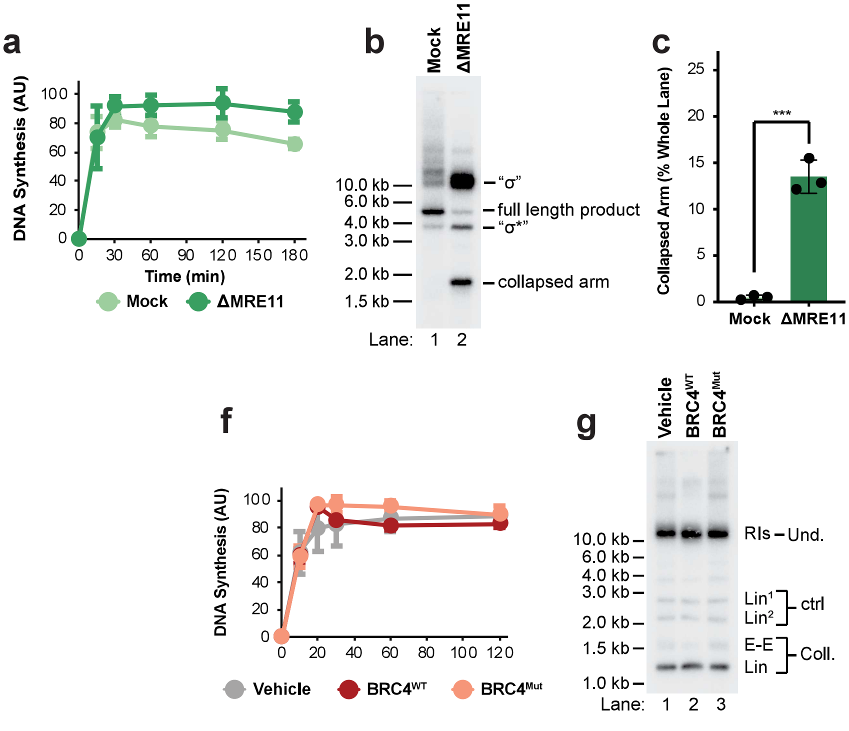

#### Supp Figure 5

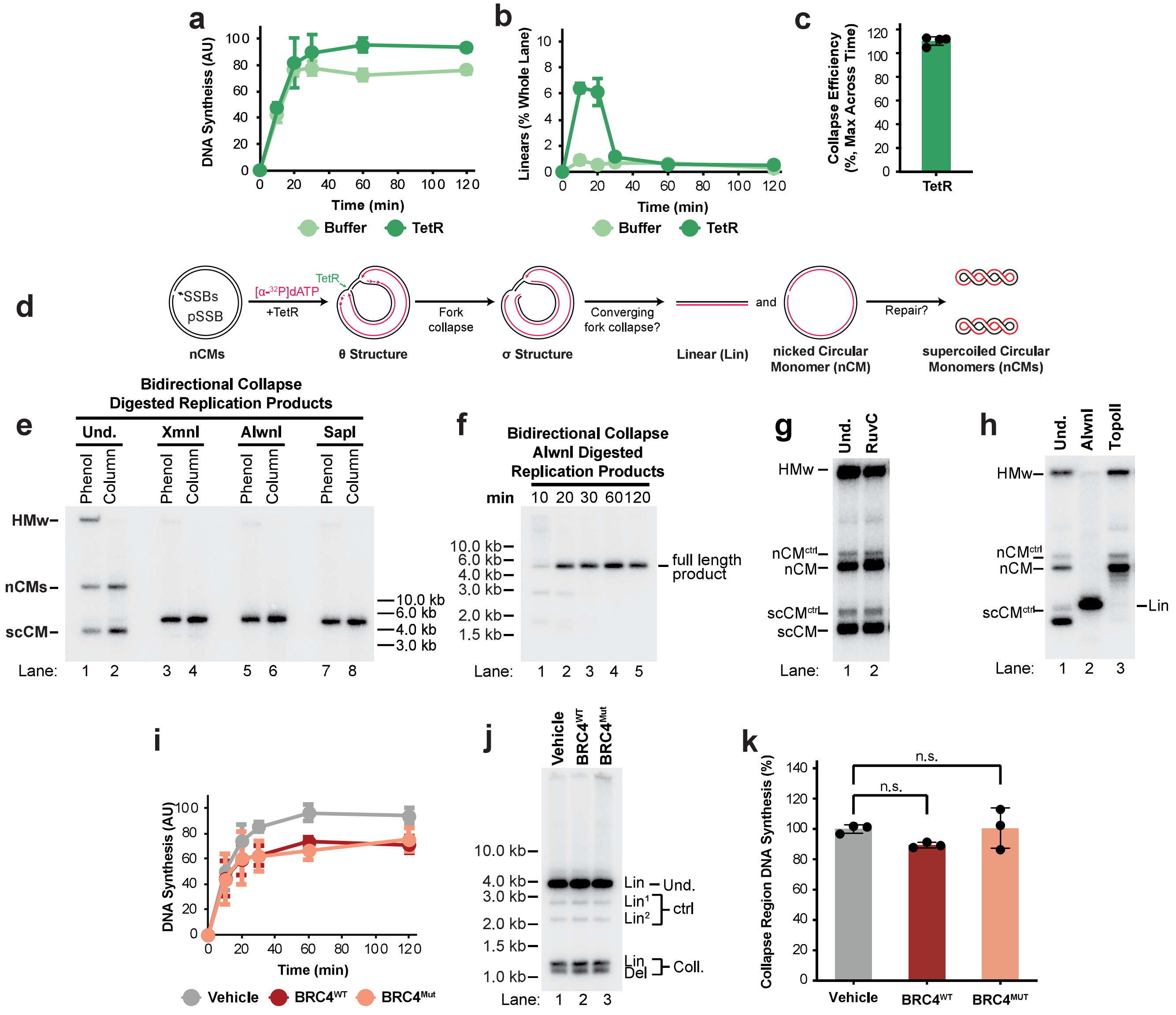

#### Supp Figure 6

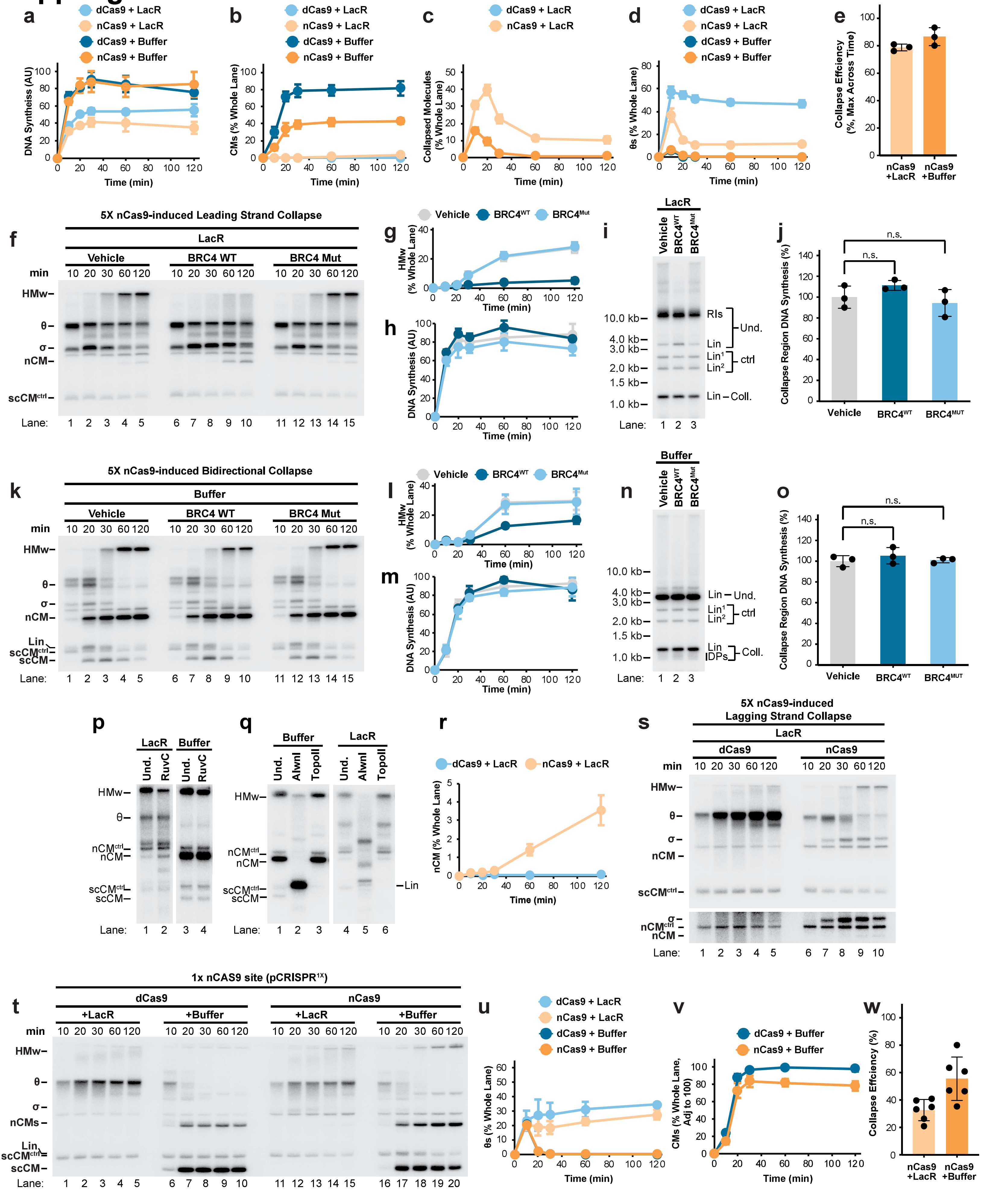

#### Supp Figure 7

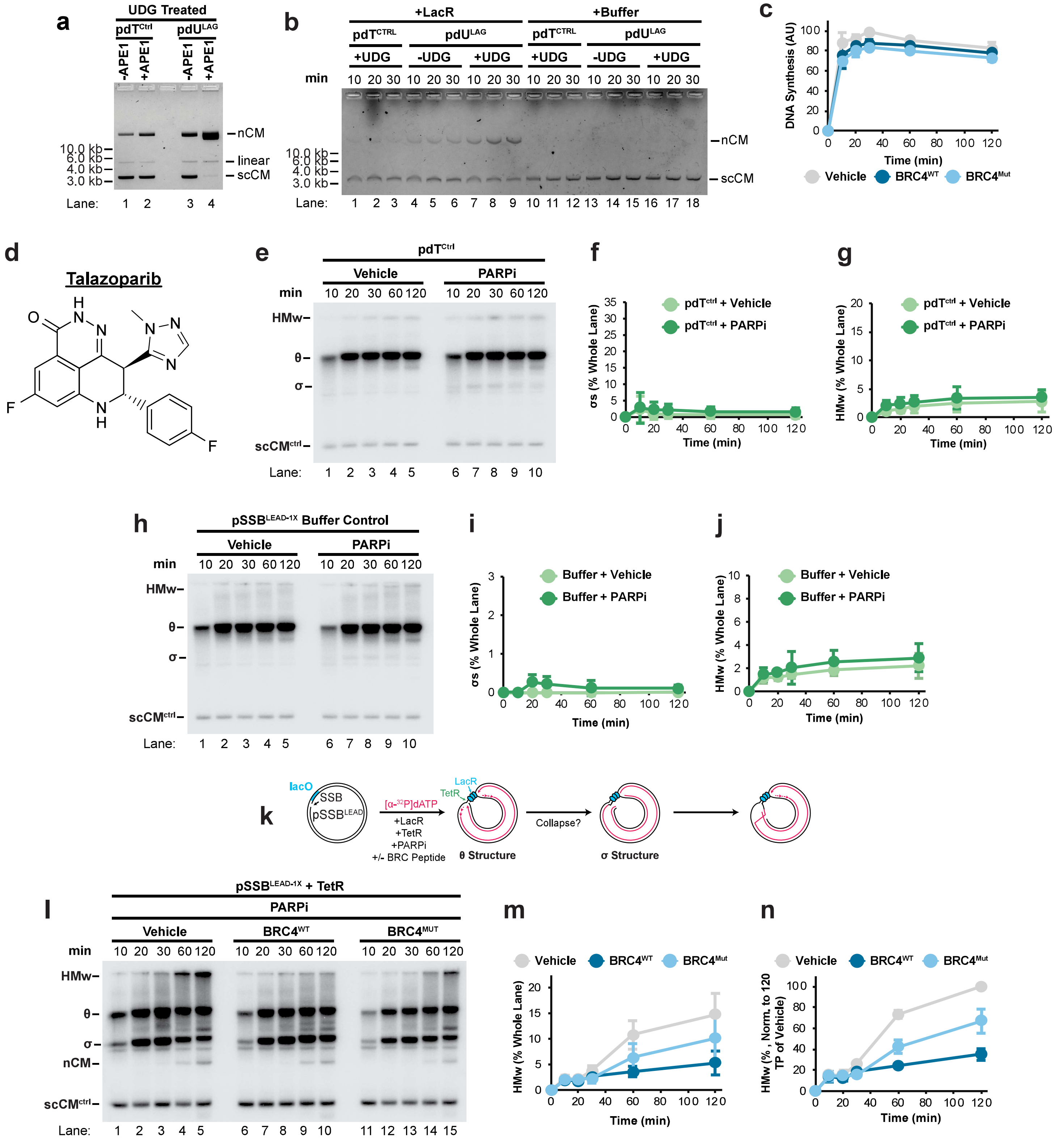
